# Structure of the chromatin remodelling enzyme Chd1 bound to a ubiquitinylated nucleosome

**DOI:** 10.1101/290874

**Authors:** Ramasubramanian Sundaramoorthy, Amanda L. Hughes, Hassane El-Mkami, David Norman, Tom Owen-Hughes

## Abstract

ATP-dependent chromatin remodelling proteins represent a diverse family of proteins that share ATPase domains that are adapted to regulate protein-DNA interactions. Here we present structures of the yeast Chd1 protein engaged with nucleosomes in the presence of the transition state mimic ADP-beryllium fluoride. The path of DNA strands through the ATPase domains indicates the presence of contacts conserved with single strand translocases and additional contacts with both strands that are unique to Snf2 related proteins. The structure provides connectivity between rearrangement of ATPase lobes to a closed, nucleotide bound state and the sensing of linker DNA. Two turns of linker DNA are prised off the surface of the histone octamer as a result of Chd1 binding, and both the histone H3 tail and ubiquitin conjugated to lysine 120 are re-orientated towards the unravelled DNA. This indicates how changes to nucleosome structure can alter the way in which histone epitopes are presented.

## Introduction

The extended family of ATPases related to the yeast Snf2 protein act to alter DNA protein interactions (Flaus *et al.*, 2006; Narlikar *et al.*, 2013). They act on a diverse range of substrates. For example, while the Mot1 protein acts on complexes between the TATA box binding protein BP and DNA (*Wollmann et al., 2011*), the Snf2 protein carries out ATP dependent nucleosome disruption (Cote *et al.*, 1994). At the heart of all these proteins are paired domains capable of rearranging during the ATP hydrolysis cycle to create a ratchet like motion along DNA in single base increments (Clapier *et al.*, 2017; Gu & Rice, 2010; Velankar *et al.*, 1999).

The yeast Chd1 protein is a member of this protein family and acts to organise nucleosomes over coding regions (Gkikopoulos *et al.*, 2011; Ocampo *et al.*, 2016; Pointner *et al.*, 2012; Tran *et al.*, 2000). Consistent with this Chd1 is known to interact with elongation factors including the Spt4-Spt5 proteins, Paf1 and FACT (Kelley *et al.*, 1999; Krogan *et al.*, 2002; Simic *et al.*, 2003).The partially redundant functions of Chd1 and Isw1 in organising nucleosomes over coding regions are in turn required to prevent histone exchange and non-coding transcription (Hennig *et al.*, 2012; Radman-Livaja *et al.*, 2012; Smolle *et al.*, 2012).

In addition to the positioning of nucleosomes, the distribution of many histone modifications is ordered with respect to promoters (Liu *et al.*, 2005; *Mayer et al., 2010*). For example histone H3 K4 methylation is frequently observed at promoters, while histone H3 K79 and K36 trimethylation are detected in coding regions (Kizer *et al.*, 2005; Li *et al.*, 2003; Pokholok *et al.*, 2005). Histone H2B is also observed to be ubiquitinylated within coding regions (Fleming *et al.*, 2008; Xiao *et al.*, 2005). Ubiquitinylation of histone H2B at lysine 123 in budding yeast, H2B K120 (H2BK120ub) in mammals, is dependent on the E2 ligase Rad6 (Robzyk *et al.*, 2000) and the E3 ligase Bre1 (Hwang *et al.*, 2003; Wood *et al.*, 2003) and removed by the deubiquitinases Ubp8 and Ubp10 (Bonnet *et al.*, 2014; Schulze *et al.*, 2011; Wyce *et al.*, 2007). A specific reader of H2BK120ub has not been identified. However, H2BK120ub does assist the histone chaperone FACT in enabling transcription through chromatin (Pavri *et al.*, 2006), and has been found to be required for methylation of histone H3 K4 and K79 (Sun & Allis, 2002). An intriguing aspect of H2BK120ub is that while mutation of the writer enzymes or K120 itself disrupts nucleosome organisation, deletion of the deubiquitinylases increases chromatin organisation (Batta *et al.*, 2011). One way in which H2BK120 may influence nucleosome organisation is via effects on enzymes responsible for chromatin organisation. Consistent with this H2BK120 increases nucleosome repositioning mediated by Chd1 (Levendosky *et al.*, 2016).

Yeast Chd1 serves as a useful paradigm in that it functions predominantly as a single polypetide. In addition, the catalytic core of the enzyme has been crystallised in association with the adjacent tandem chromodomains (*Hauk et al., 2010*). Similarly the C-terminal region of the protein has been crystalized revealing that this region includes SANT and SLIDE domains that comprise the DNA binding domain (DNABD) (Ryan *et al.*, 2011b; Sharma *et al.*, 2011) and are also present in ISWI proteins (Grune *et al.*, 2003). Chd1 enzyme engages nucleosomes in a conformation in which the SANT and SLIDE domains bind linker DNA, while the ATPase domains engage DNA at super helical location (SHL) 2 (Nodelman *et al.*, 2017; Sundaramoorthy *et al.*, 2017). Higher resolution structures of both Chd1 (Farnung *et al.*, 2017) and Snf2 (Liu *et al.*, 2017) show that the ATPase domains make contacts with DNA via residues that are conserved in ancestral single stranded ATPases and some unique to Snf2 related ATPases. The binding of the Chd1 DNABD unravels two turns of DNA from the surface of nucleosomes in a nucleotide stimulated reaction (Farnung *et al.*, 2017; Sundaramoorthy *et al.*, 2017). Here we report a structure for the yeast Chd1 protein in association with a nucleosome, bearing modifications that are found to occur within coding regions, where Chd1 is known to act. Interestingly, nucleosomal epitopes are observed to be reconfigured specifically on the side of the nucleosome on which DNA is unwrapped. This indicates the potential for changes to nucleosome structure to reconfigure the way in which histone epitopes are presented.

## Results

### The structure of Chd1 nucleosome complexes

As Chd1 functions on transcribed genes, it is of interest to understand the interplay between Chd1 and histone modifications observed in coding region chromatin. As a result nucleosomes were prepared in which histone H3 K36 was alkylated to mimic trimethylation (Figure 1 – Figure supplement 1) and H2B cross-linked to ubiquitin (Figure 1 – Figure supplement 2). Conditions were established to favour binding of a single Chd1 to modified nucleosomes that included an asymmetric linker DNA extension of 14bp (Figure 1 – figure supplement 2B) in the presence of ADP-BeF. Purified complexes were frozen onto EM grids.

**Figure 1.**
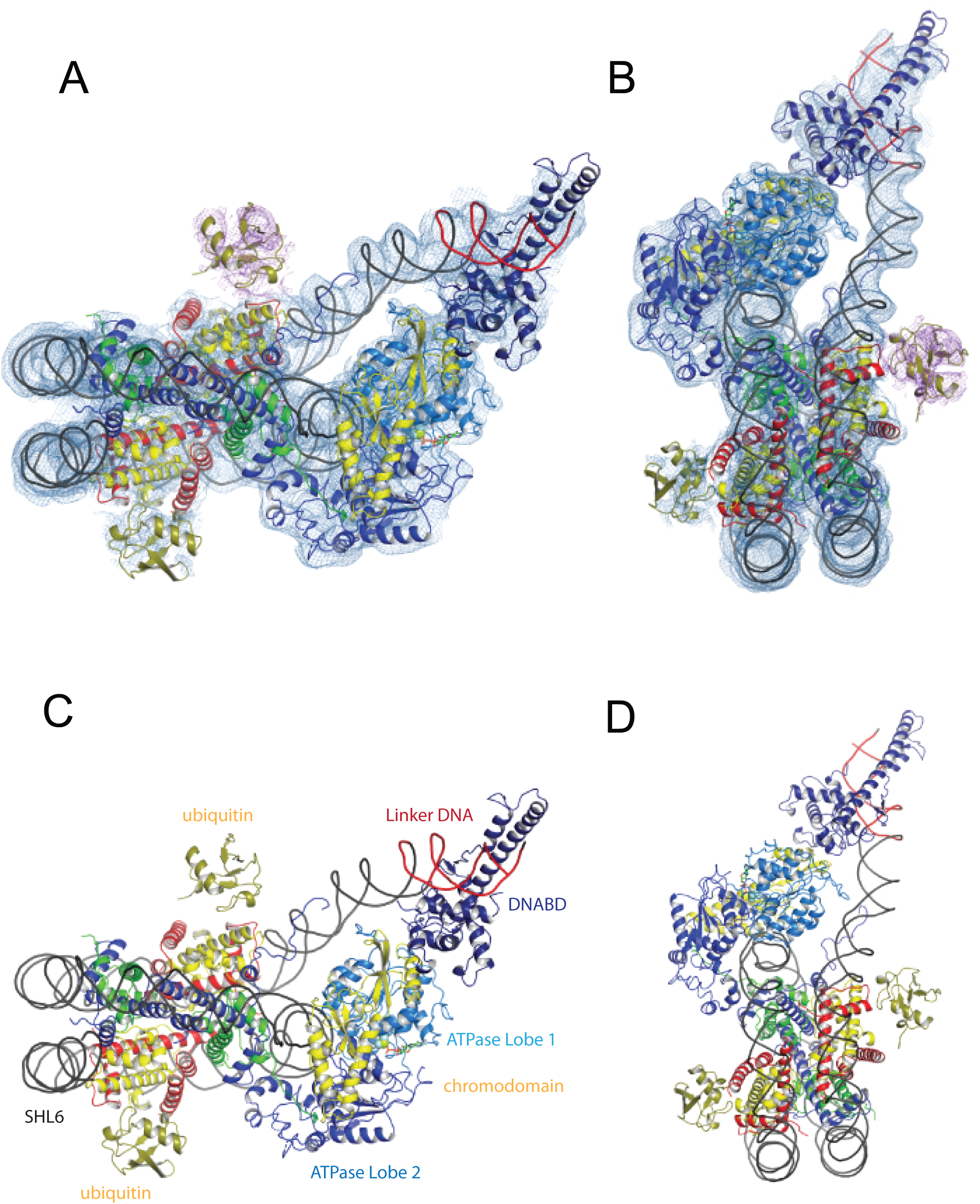
A Chd1-Nucleosome complex. (**A**, **B**) Overall fit of nucleosome bound Chd1 to density map. Chd1 chromodomains – yellow, DNABD – dark blue, ATPase lobe 1 cyan, ATPase lobe 2 blue, Ubiquitin dark yellow, H2B yellow, H2A red, H3 green, H4 blue. (**C**, **D**) two views of the structural model.

**Figure 1 – Figure supplement 1.**
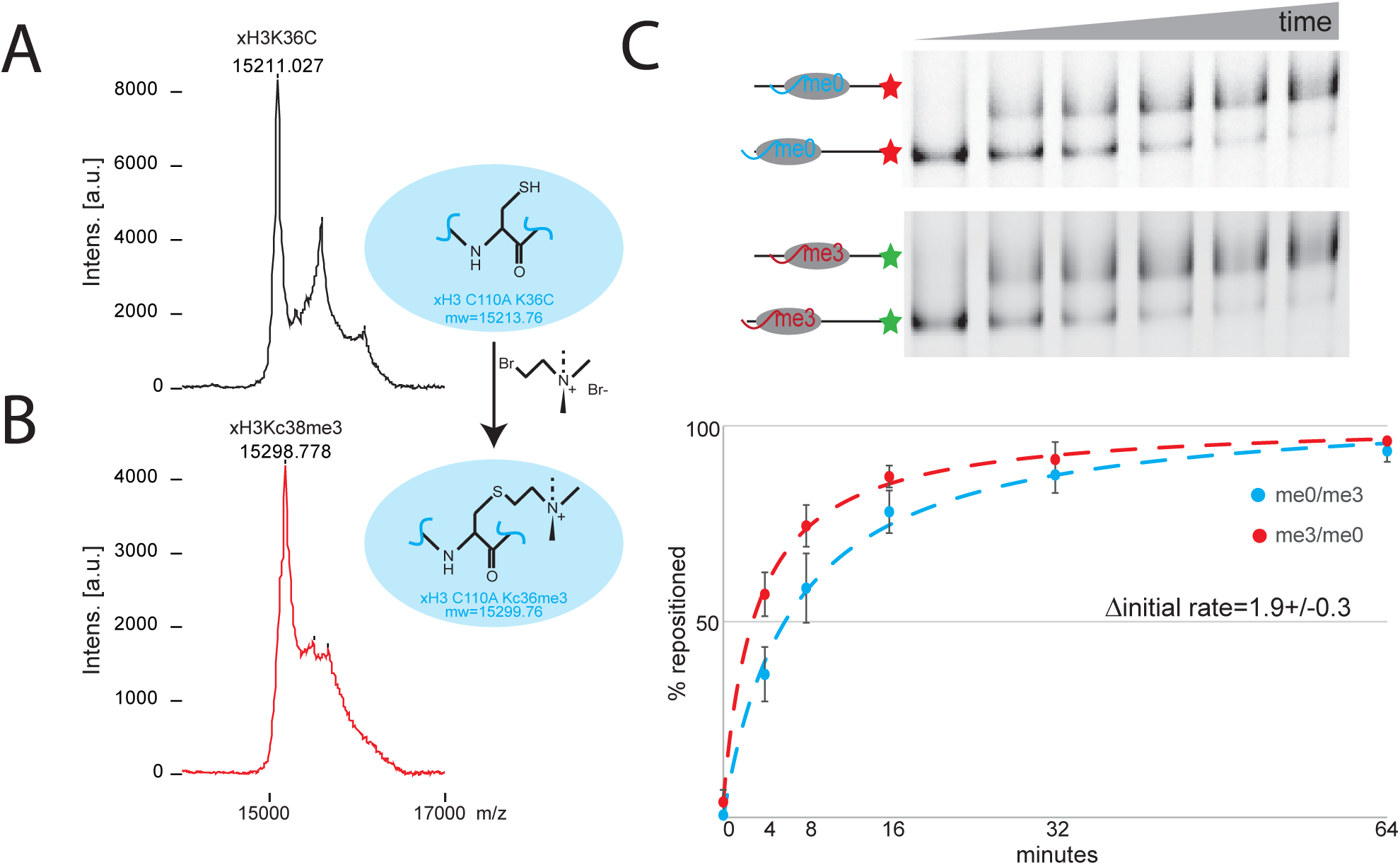
Preparation of nucleosomes with an alkylation mimic of histone H3 K36 methylation. *Xenopus laevis* histone H3K36C was alkylated to generate a mimic of trimethyl lysine (Simon *et al.*, 2007). MALDI mass spectrometry shows that the mass of free histone H3 (**A**) increases by 86Da following alkylation (**B**) consistent with efficient conversion. (**C**) Chd1 directed nucleosome repositioning was assessed using a two colour nucleosome repositioning assay in which differently modified octamers are assembled onto DNA fragments labelled with different fluorescent dyes assays. Comparison of repositioning rates from 4 repeats indicates that the initial rate of repositioning is 1.9 fold greater for the trimethyl mimic.

**Figure 1 – Figure supplement 2.**
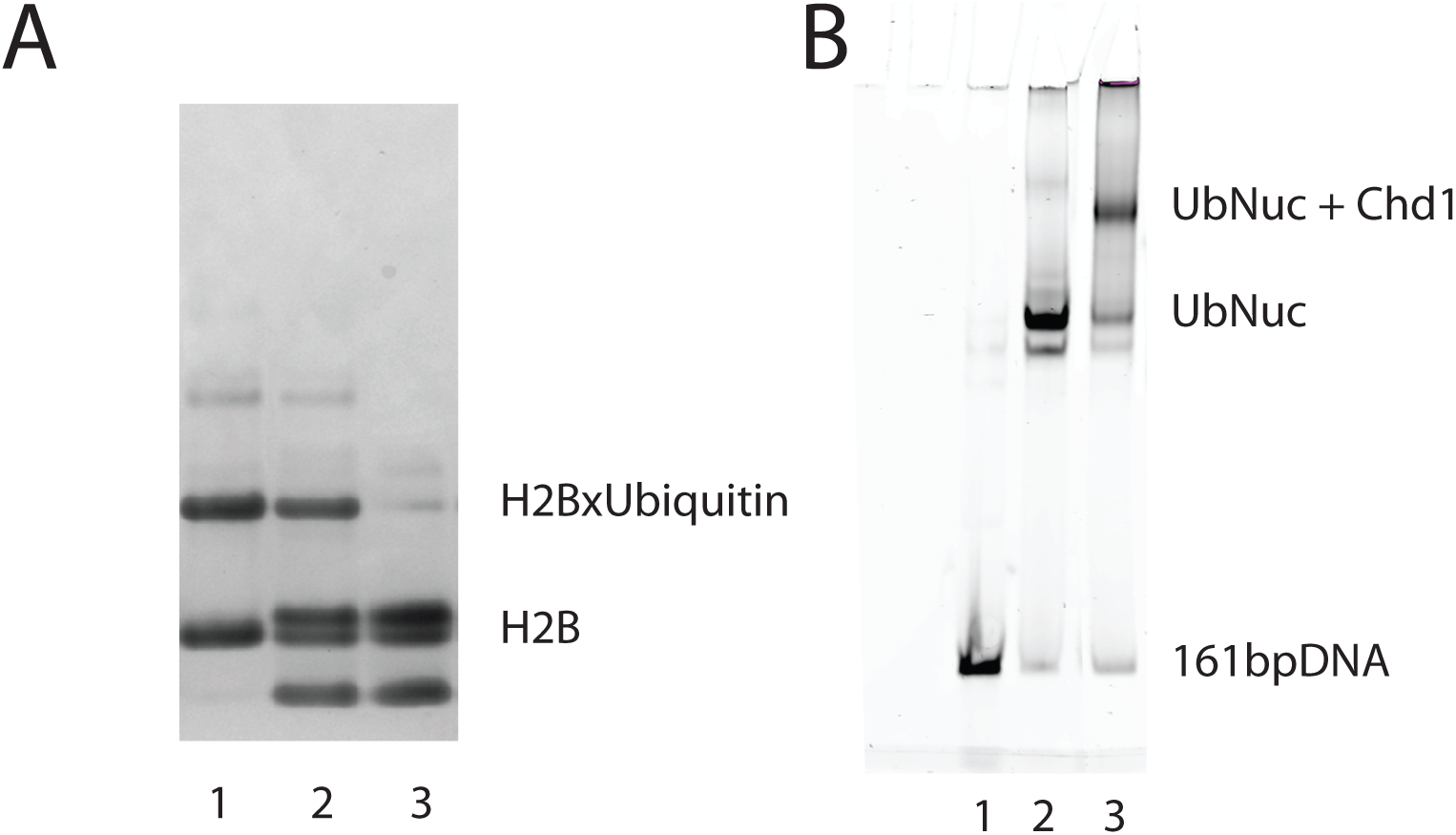
Generation of nucleosomes with ubiquitin cross-linked to histone H2B K120. Ubiquitin was cross-linked to H2BK120C as described previously (Long *et al.*, 2014; Morgan *et al.*, 2016). Ubiquitin modified H2B was purified by ion exchange chromatography and the extent of coupling confirmed by SDS PAGE (**A**). Nucleosomes were assembled on DNA including an asymmetric 14 base pair linker using ubiquitin modified H2B and alkylated H3K36. (**B**) The assembly of nucleosomes was assessed by native PAGE (lane 2). Incubation with Chd1 resulted in a major species consistent with binding of a single Chd1 to a nucleosome (lane 3).

2D classification of some 893000 particles revealed 16 classes in which nucleosomes with the Chd1 molecule attached could be identified (Figure 1 – Figure supplement 3B). Initial 3D classification resulted in 5 related classes (Figure 1 - Figure supplement 3C). Three of these were combined and reclassified as six classes, one of which was selected for refinement. This resulted in the generation of a map with a resolution of 4.5Å (FSC 0.143) (Figure 1 – Figure supplement 4A). The resolution varies within the map, with resolution close to 4 Å in the region occupied by the nucleosome and ATPase lobes and lower resolution in the vicinity of the DNABD and ubiquitin peptides (Figure 1 – figure supplement 4B). The nucleosome particles exhibited a preferred orientation which may limit the resolution (Figure 1 Figure supplement 4C). A structural model was generated to fit the density map making use of the structures of a nucleosome assembled on the 601 DNA sequence, Chd1 chromoATPase, and DNABD (Figure 1). The fit for individual components of the structure to the electron density is shown in Figure 1 - Figure supplement 5.

**Figure 1 – Figure supplement 3.**
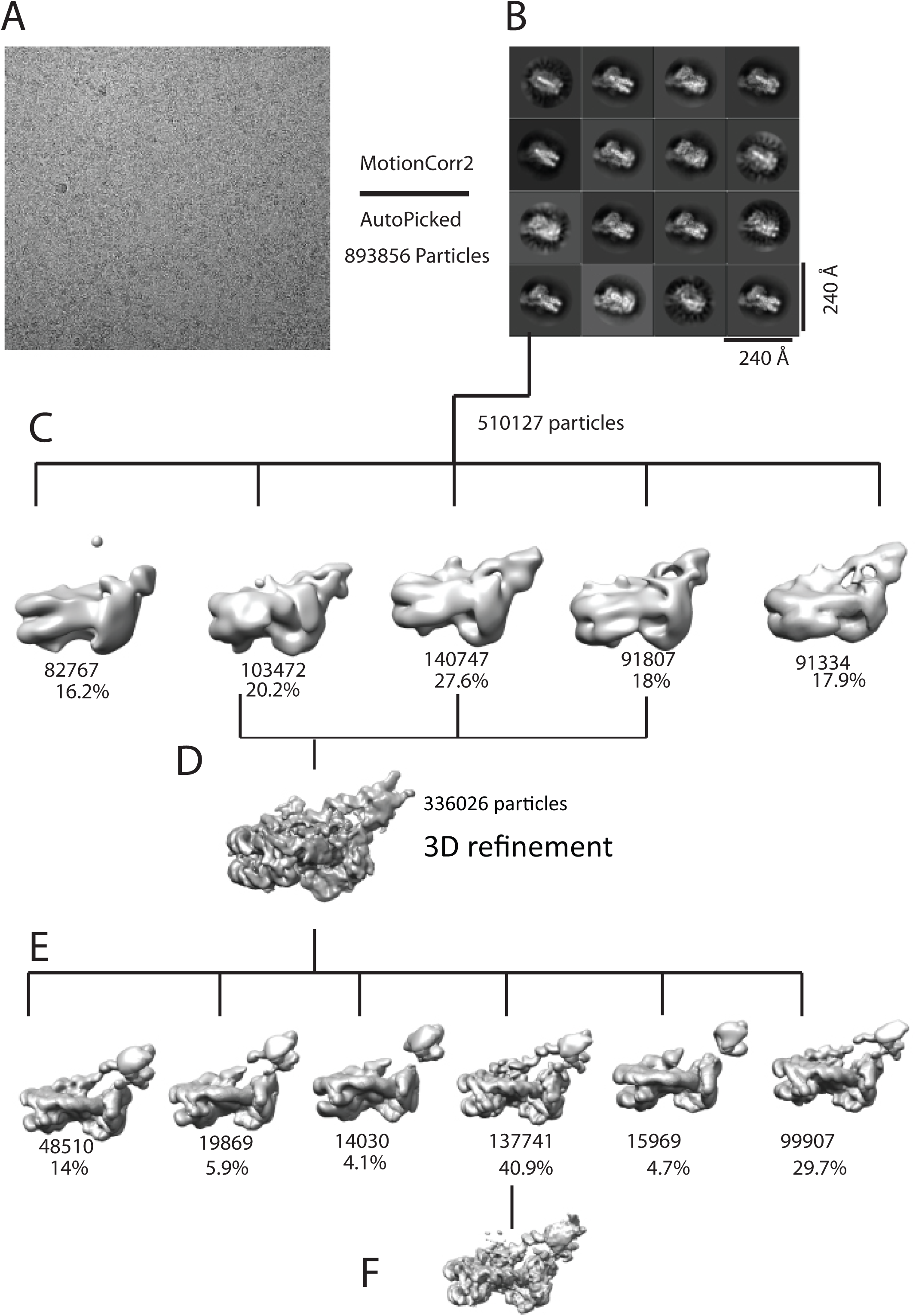
Image processing for Chd1-nuclesome complex. **A)** An example motion corrected micrograph. **B)** 2D classification resulted in identification of 16 classes with Chd1 bound to nucleosomes. **C)** Initial 3D classification resulted in five classes three of which were combined **D**) reclassified **E**) and refined **F**).

**Figure 1 – Figure supplement 4.**
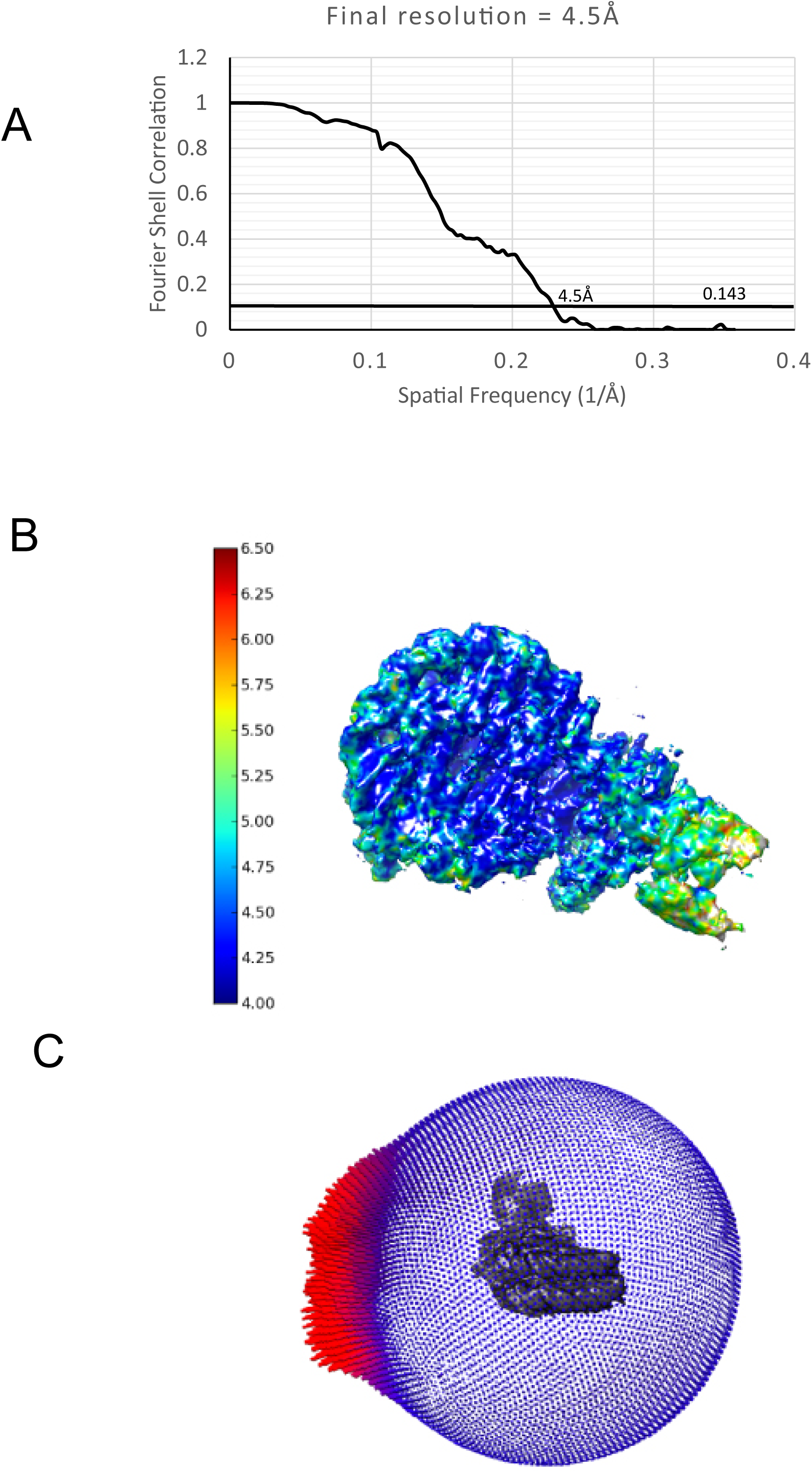
Properties of the Chd1-nuclesome complex. **(A)**Average resolution was estimated from a plot of the fourier shell correlation between the two half maps. (**B**) Plot showing local distribution of resolution. (**C**) Angular distribution of particles used for final refinement.

**Figure 1 – Figure supplement 5.**
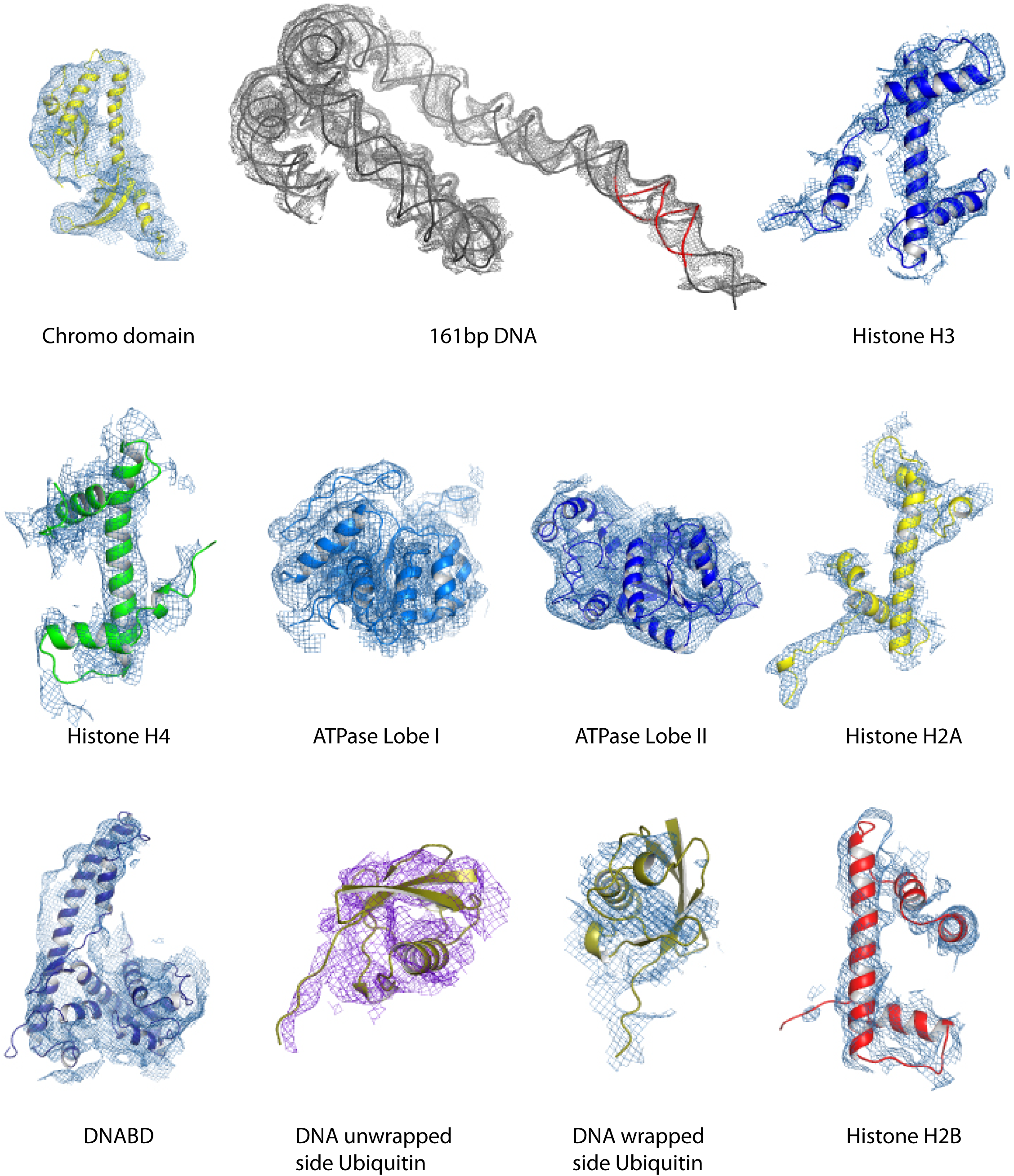
Fit of individual components. The indicated components fitted individually to relevant regions of the density map.

The overall organisation of Chd1 is similar to that observed previously by cryo EM (Farnung *et al.*, 2017; Sundaramoorthy *et al.*, 2017) and directed cross-linking (Nodelman *et al.*, 2017). The ATPase domains are bound at the SHL-2 location. Of the two SHL2 locations within nucleosomes, the bound site is in closest proximity to SANT-SLIDE domain bound linker DNA in physical space, but distal on the unwrapped linear DNA sequence (Figure 1). Chd1 predominantly contacts the nucleosome via contacts with DNA, via the DNABD in the linker and ATPase lobes at SHL2, contacts with histones are limited to the histone H3 and H4 N-terminal regions discussed below.

We previously showed that Chd1 binding results in nucleotide-dependent unwrapping of nucleosomal DNA resulting from the interaction of the DNABD with linker DNA (Sundaramoorthy *et al.*, 2017). The higher resolution of the current structure shows that precisely two turns of nucleosomal DNA are unravelled (Figure 1). The extent of DNA unwrapping observed here when Chd1 is bound to nucleosomes flanked by a 14 base pair linker DNA is identical to that observed when Chd1 is bound to the opposite surface of the 601 nucleosome positioning sequence with a 63 base pair linker (*Farnung et al., 2017*). As the interaction of histones with the two sides of the 601 positioning sequence differ quite dramatically (Chua *et al.*, 2012; Hall *et al.*, 2009; Levendosky *et al.*, 2016; Ngo *et al.*, 2015), this suggests that the extent of unwrapping is dominated by the properties of Chd1 rather than the affinity of DNA for the octamer. The path of this unwrapped DNA is oriented away from the plane of the wrapped DNA gyre and is kinked at the location where contacts are made with the SANT-SLIDE domains (Figure 1). Other than DNA unwrapping we do not detect additional changes in the organisation of DNA on Chd1 bound nucleosomes at this resolution.

The orientation of the DNABD is critical in determining the extent of DNA unwrapping. The only contacts detected between the DNABD and the remainder of Chd1 are contacts with the chromodomains (Figure 2). The first of these is the interaction between K329 of chromodomain II and D1201 P1202 in the SLIDE component of the DNABD and has been observed previously (Farnung *et al.*, 2017; Nodelman *et al.*, 2017)(Figure 2 – Figure supplement 1A). The second contact is between S344 and K345 in the linker helix between chromodomain II and ATPase lobe I with the SANT component of the DNA binding domain at D1033-D1038 (Figure 2 – Figure supplement 1A). Given that chromodomains are present in Chd1 enzymes but not ISWI and Snf2 remodellers it makes sense that the residues contacted in the SANT and SLIDE domains are most highly conserved in Chd1 proteins (Figure 2 – Figure supplement 1B). The interaction of the DNABD with linker DNA prises off two turns of nucleosomal DNA, a process that likely requires substantial force (Hall *et al.*, 2009; Meng *et al.*, 2015). As a result it is somewhat surprising that the area of contact between the chromodomains and DNABD is so small. It is possible that additional regions of Chd1 including the N-terminus that is not resolved in the density map also contribute to this interaction as suggested by previous studies of Chd1 proteins (Liu *et al.*, 2015; Zhou *et al.*, 2018)(Sundaramoorthy *et al.*, 2017).

**Figure 2.**
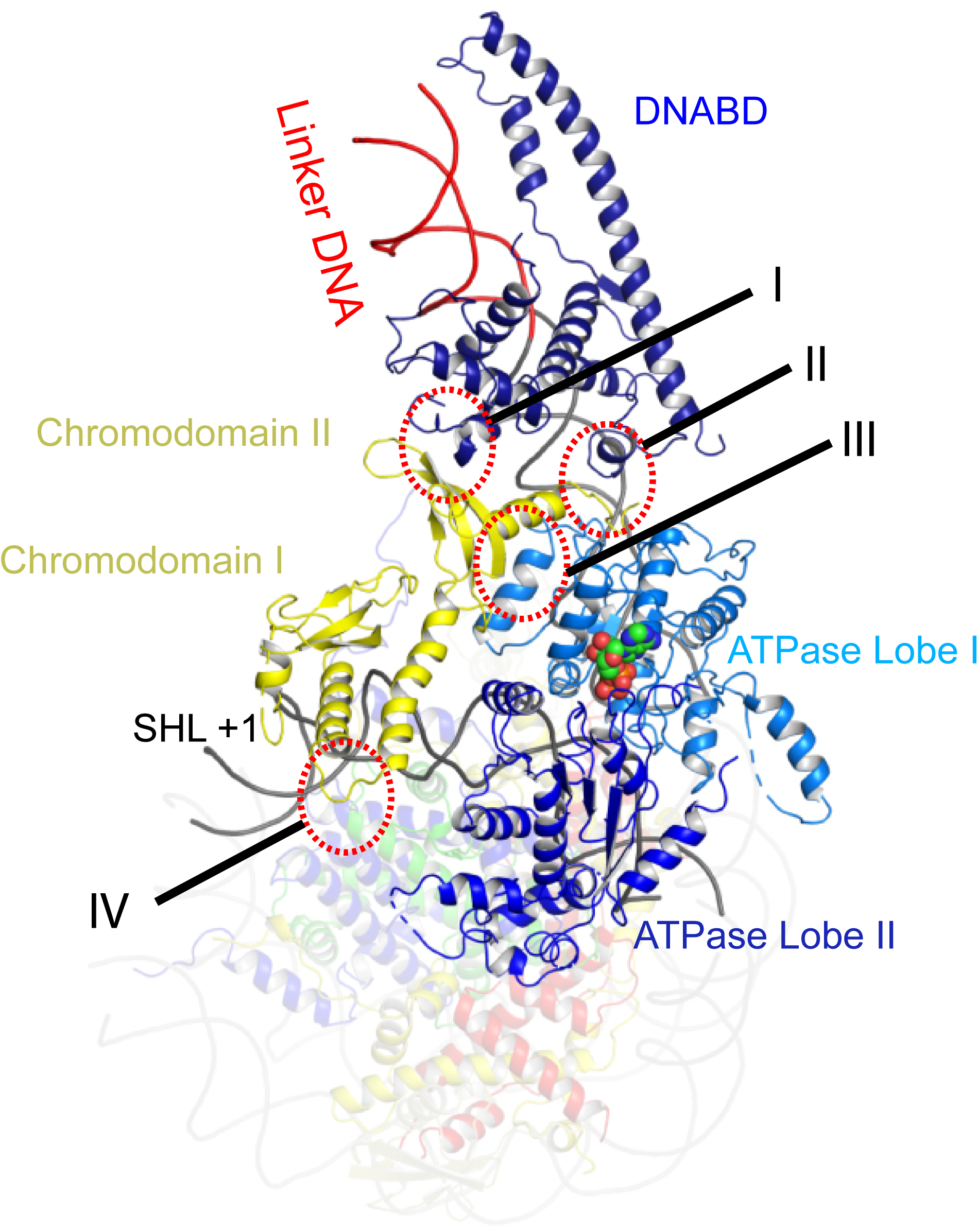
Contacts constraining Chd1 chromodomains. Overview of the major contacts constraining the positioning of the Chd1 chromodomains. Colours of domains as for Figure 1. Key contacts are highlighted. I) Chromodomain II SLIDE, II) Chromodomain linker helix to SANT, III) Chromodomain II to ATPase lobe 1, IV) Chromodomain I to nucleosomal DNA at SHL +1.

**Figure 2 – Figure supplement 1.**
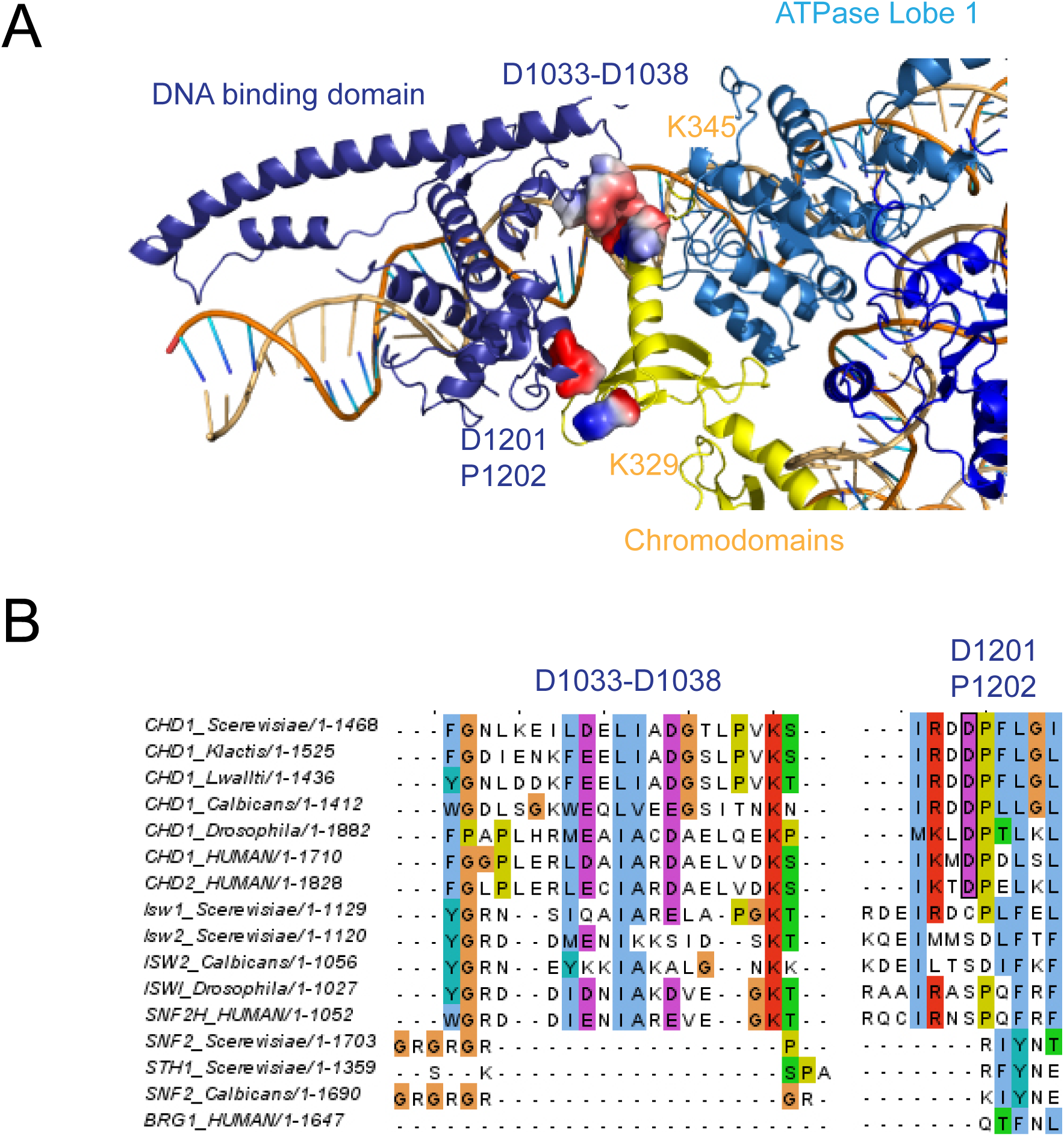
The chromodomain contacts with SANT and SLIDE domains. **A)** Residues involved in the interactions are coloured by surface charge. **B)** Sequence alignments showing conservation of residues making these contacts in Chd1 proteins.

### Repositioning of Chd1 ATPase lobes to a closed ATP-bound state drives repositioning of chromodomains

The position of chromodomains is determined by each of the four contacts made with other components of the complex (Figure 2). When not bound to nucleosomes, the tandem chromodomains of Chd1 are observed to impede DNA binding to the ATPase domains (Hauk *et al.*, 2010). This gave rise to the prediction that these domains would be rearranged in the nucleosome bound state (Hauk *et al.*, 2010). This is indeed the case as the chromodomains undergo an 18 degree rotation when compared to the orientation observed in the crystal structure of Chd1 in the open state (Figure 3). Following repositioning chromodomain I interacts with nucleosomal DNA at SHL1 (Figure 2 – Figure supplement 2) as observed previously (Farnung *et al.*, 2017; Nodelman *et al.*, 2017).

**Figure 3.**
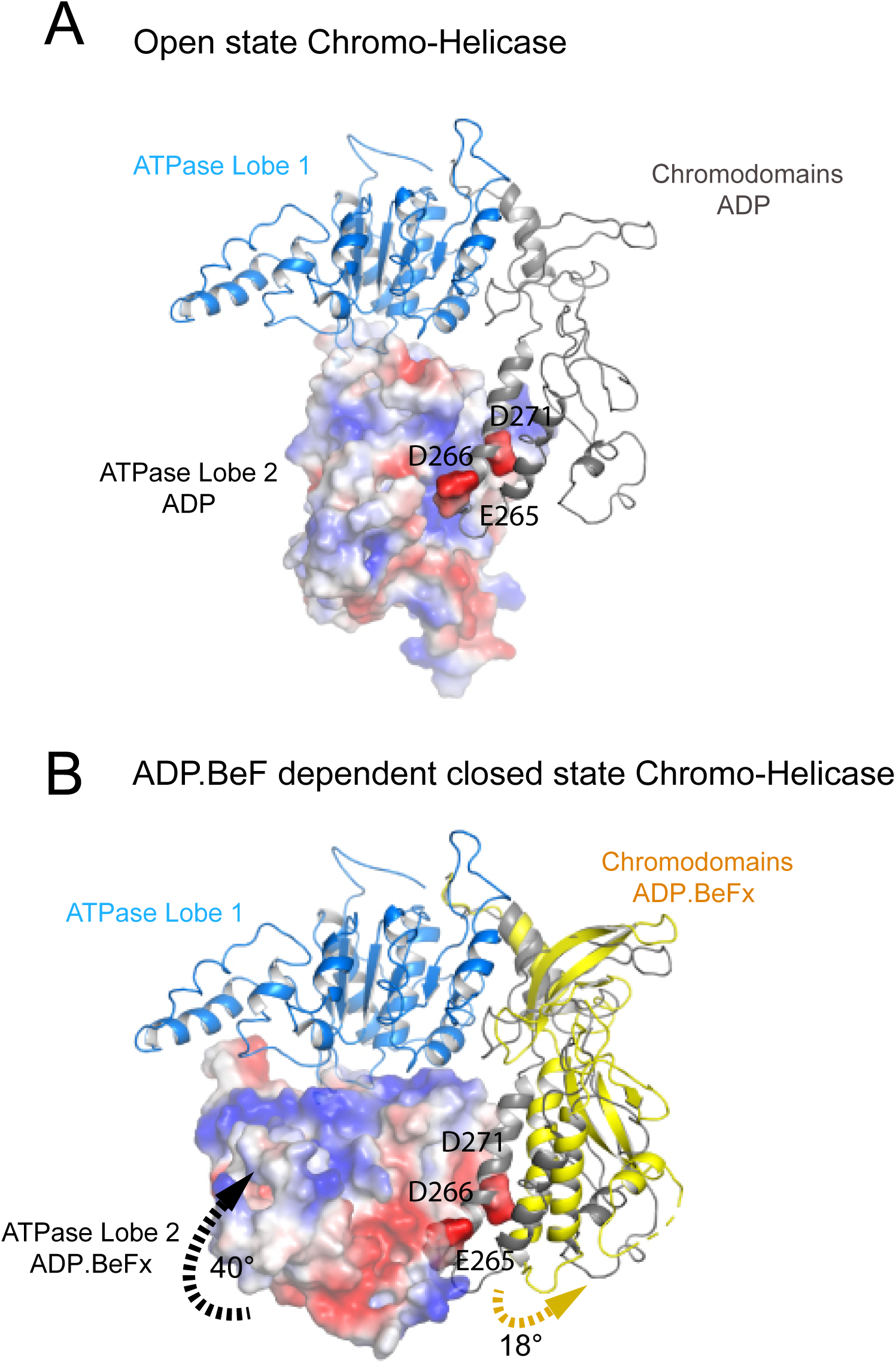
Closure of the ATPase lobes changes the chromodomain interaction surface. **A**) The long acidic helix within chromodomain I interacts with a basic surface on ATPase lobe 2 in the open state (3MWY). **B)** In the closed state the basic surface on lobe 2 is rotated towards DNA and replaced with an acidic region. The long acidic helix within chromodomain I is repositioned away from this acidic surface.

**Figure 2 – Figure supplement 2.**
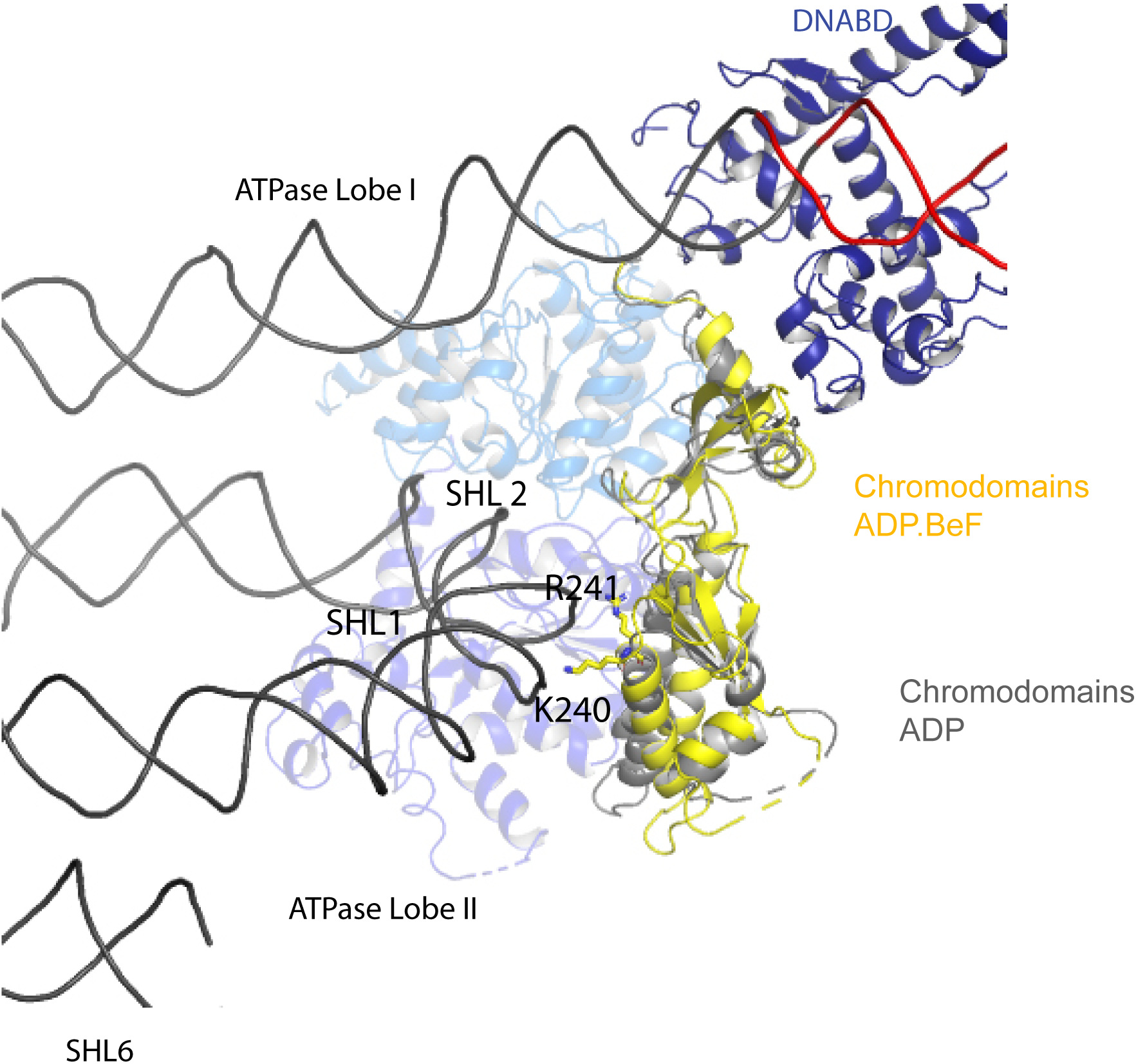
Chromodomain interactions with nucleosomal DNA at SHL+1. Interaction of Chd1 chromodomains with nucleosomal DNA at SHL +1. R241 and K240 come in close contact with DNA.

Coincident with repositioning of the chromodomains, ATPase lobe II is repositioned closer to lobe I. This results in residues including those contributing to the conserved Walker box motifs (K407 and R804, R807) being brought into an arrangement compatible with ATP catalysis. Density for ADP-BeF within the pocket formed by conserved residues from ATPase domains I and II is well defined (Figure 2 – Figure supplement 3).

**Figure 2 – Figure supplement 3.**
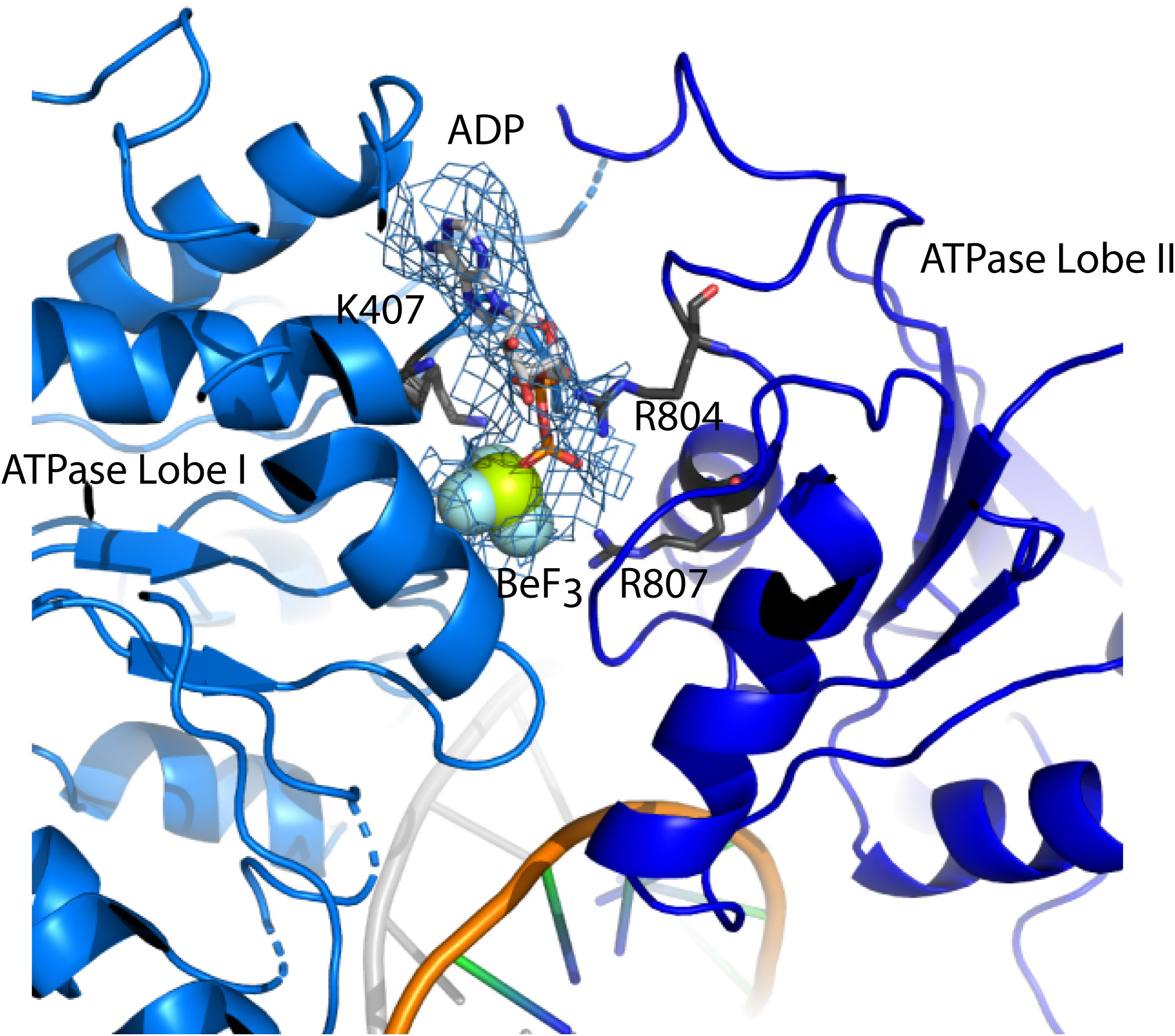
The ATP binding site. ADP Berylium Fluoride fitted to density in a pocket formed by conserved residues contributed by both ATPase domains which are positioned in a closed state. Residues observed to contribute to catalysis in related proteins are well positioned to function similarly in Chd1.

The repositioning of ATPase lobe II enables contacts to be made with nucleosomal DNA (see below), the histone H4 tail and the histone H3 alpha 1 helix (Figure 2 – figure supplement 4).These are the only direct contacts with histone components of the nucleosome. The contact with the H4 tails is conserved in mtISWI and Snf2 (Liu *et al.*, 2017; Yan *et al.*, 2016). D729 and E669 are conserved across all classes of remodelling enzyme but D725 is not as well conserved in Snf2 related enzymes (Figure 2 – figure supplement 4B). The conservation of this contact in Chd1 enzymes is consistent with the H4 tail playing an important role in regulating Chd1 activity; deletion or mutation of the H4 tail has been shown to reduce nucleosome sliding and ATPase activity (Ferreira *et al.*, 2007). The additional helices that make up the protrusion 2 region of ATPase lobe 2 in Chd1 are conserved in chromatin remodeling ATPases, but not within all SF2 DNA translocases. Within this region residues 638-642 interact with the alpha 1 helix of histone H3 (Figure 2 – figure supplement 4A). The residues participating in the interaction are progressively less well conserved in ISWI and SNF2 related proteins (Figure 2 – figure supplement 4C). A loop from Phe1033 to Leu 1045 in an equivalent region of the yeast Snf2 protein is not assigned in the Snf2-nucleosome structure, but this region is positioned such that a related contact with histone H3 could be made.

The structure, also provides clues as to how these conformational changes are driven. A central event is likely to be the closure of the cleft between ATPase domains driven by ATP binding (Figure 2 – Figure supplement 3). The 40° rotation of ATPase lobe II required to form the ATP binding pocket results in a negatively charged surface, observed to interact with an acidic surface on the long helix of chromodomain I (Figure 3A)(Hauk *et al.*, 2010), being replaced by an acidic surface likely to repel chromodomain I (Figure 3B). As a result closure of the ATPase domains is anticipated to drive nucleotide dependent repositioning of the chromodomains. Pulsed EPR was used to directly measure repositioning of the chromodomains in the absence of nucleosomes (Figure 4). The distance between engineered label at V256C in chromodomain I and S524C in ATPase lobe1 is 4.4nm in the open state, consistent with that observed in the crystal structure of the Chd1 chromoATPase domains (Hauk *et al.*, 2010). In the presence of ADP-BeF the 4.4nm distance predominates, but a shoulder is observed consistent with a proportion of molecules adopting a new conformation with a distance of 5.6nm (Figure 4) which is similar to that observed in the ADP-BeF bound nucleosome by cryo-EM. This indicates that ATP binding is a driving event for repositioning of the chromodomains.

**Figure 4.**
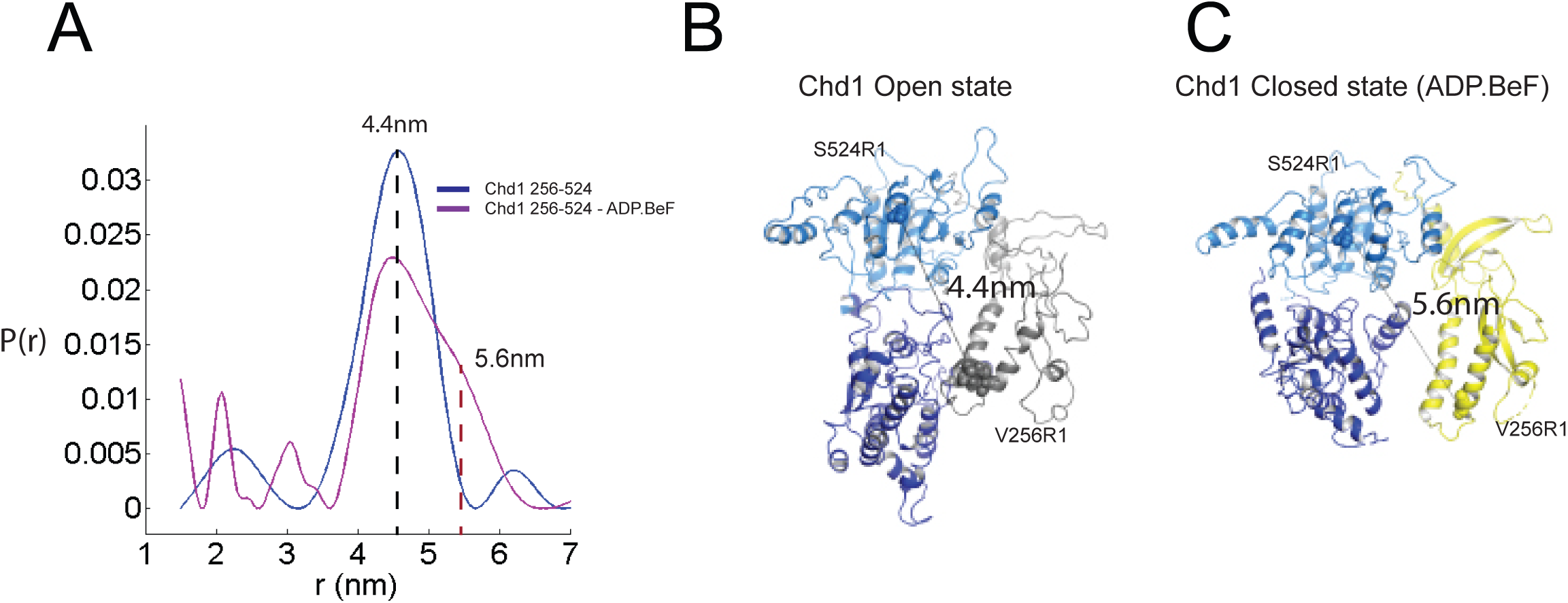
Nucleotide dependent reconfiguration of Chd1 chromodomains. Pulsed electron paramagnetic measurements were used to measure the distance between nitroxyl reporter groups attached to Chd1 at ATPase lobe 1 (S524) and chromodomain I (V256). **A**) The probability distribution P(r) at different separations was measured in the presence (purple) and absence (blue) of ADP.BeF. The distance corresponding to the major distance is shown for both measurements, for the ADP.BeF sample the distance corresponding to the shoulder is also indicated. Modelled distances between these labelling sites in the open state (3MWY), and the closed state observed in the Chd1 bound nucleosome are indicated in **B** and **C** respectively.

**Figure 2 – Figure supplement 4.**
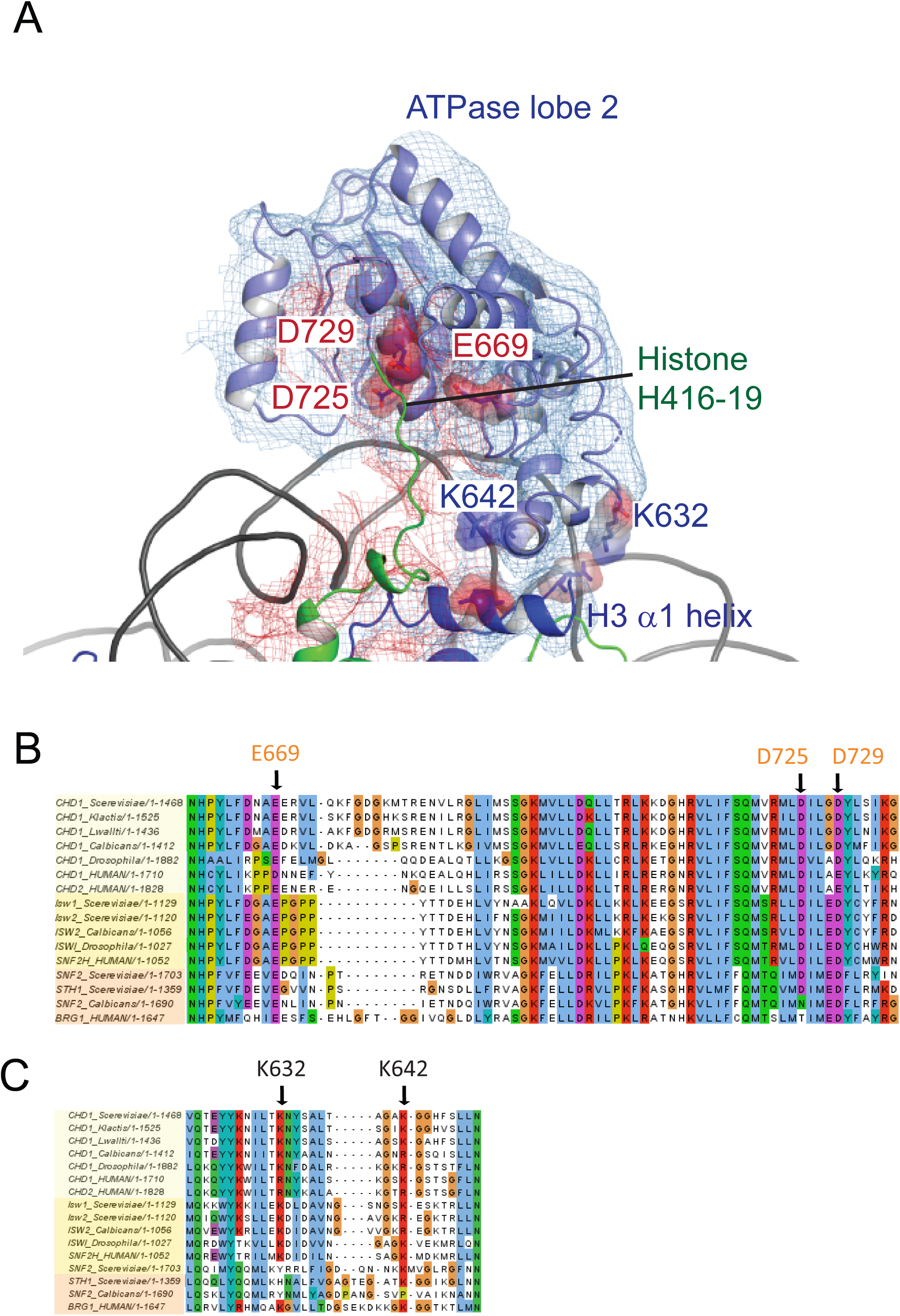
ATPase lobe 2 interaction with histone H4 tail. **A)** Fit of ATPase lobe 2, histone H3 and Histone H4 tail to density map. Residues coming into proximity with the H4 tail, D275, D279 and E669 are indicated by surface charge. The contact between ATPase lobe 2 and the histone H3 alpha 1 helix is also shown. **B** and **C,** Sequence alignments show that the sequences participating in these interactions are conserved in Chd1 proteins and to some extent in Iswi and Snf2 proteins as well.

The partial repositioning of the chromodomains observed in free Chd1 is likely to be stabilised by additional favourable interactions formed when this repositioning occurs within the context of nucleosome bound Chd1. These include the formation of contacts between chromodomain I and DNA at SHL1, between ATPase lobe II and the H3 alpha 1 helix, between ATPase lobe II and the histone H4 tail and most significantly the formation of a substantial interaction interface between ATPase lobe II and nucleosomal DNA at SHL2. The repositioning of the chromodomains in turn acts as a lever to reposition the DNA binding domain. In the context of nucleosomes this results in nucleotide dependent unwrapping of two turns of nucleosomal DNA (Sundaramoorthy *et al.*, 2017). Conversely, the interaction of the DNABD requires linker DNA to be accessible.

In order to investigate how the ability of the DNABD to interact with linker DNA is affected by the presence of an adjacent nucleosome, interactions between dinucleosomes with different separations were modelled. With a linker length of 19 bp Chd1 can be modelled binding the linker between adjacent nucleosomes (Figure 5). However, as the linker between nucleosomes is reduced, steric clashes become increasingly prohibitive. The requirement for a 19bp linker is likely to provide a limit below which engagement of the DNABD will be less stable. As this lower limit is set by clashes between the DNABD and the adjacent nucleosome, it is different from the length of linker required to occupy the DNA binding surface of the SANT and SLIDE domains on a mononucleosome with a free DNA linker. In this latter case 7 base pairs of DNA make contact with the DNABD (Figure 1). The c19 bp separation below which access of the DNABD to linker becomes progressively more difficult resonates with the average inter-nucleosome spacing of 19 bp observed in *Saccharomyces cerevisiae* (Tsankov *et al.*, 2010). As the conformation of the DNABD is connected via the chromodomains to the ATPase domains, the structure of Chd1 provides molecular connectivity between the availability of nucleosomal linker DNA in excess of 19bp and the generation of closed nucleotide bound motor domains. This potentially provides a mechanism via which linker DNA length regulates the rate of nucleosome movement (Yang *et al.*, 2006).

**Figure 5.**
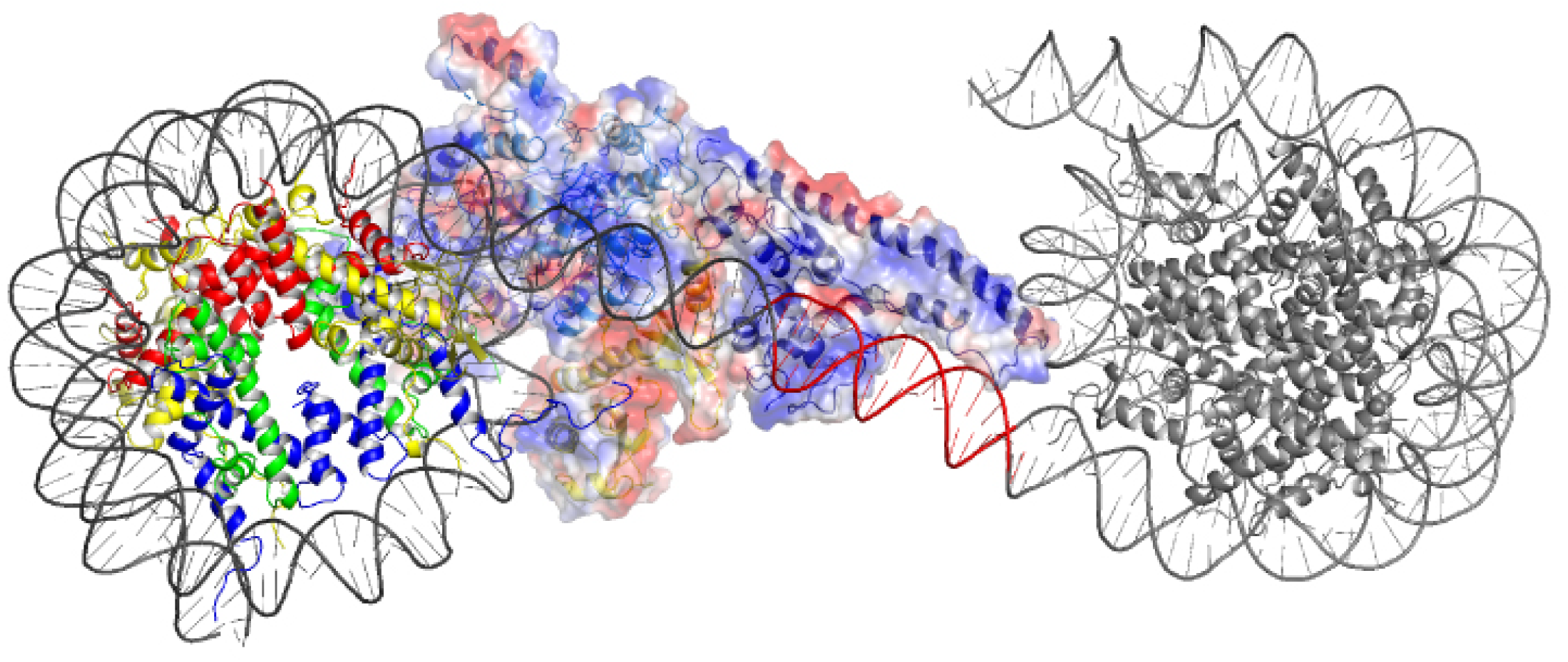
Modelling the interaction of Chd1 between adjacent nucleosomes. The structure of the Chd1 bound nucleosome was used to model binding to the linker between an adjacent nucleosome (grey) by extending the liner DNA to 19bp. With shorter linkers steric clashes with the adjacent nucleosome become progressively more severe.

### Organisation of the Chd1 ATPase domains

Nucleosome repositioning is likely to be driven by the ability of the ATPase domains to drive ATP dependent DNA translocation. This has been observed directly for several Snf2 family proteins (Deindl *et al.*, 2013; Lia *et al.*, 2006; Sirinakis *et al.*, 2011; Zhang *et al.*, 2006) and is conserved within a wider family of superfamily II ATPases (Singleton *et al.*, 2007). Structures of superfamily II single stranded translocases, such as herpes virus NS3, in different NTP bound states illustrate how the ratcheting motion of the ATPase domains drives translocation (Gu & Rice, 2010). To date such a series of structures does not exist for a double strand specific translocase. This raises the question as to whether structures of NS3 can be used to inform key aspects of the mechanism of Chd1 such as identifying the tracking strand. To do this we first align the ATPase lobes of Chd1 individually with NS3. The ATPase lobes of Chd1 like other Snf2 related proteins contain additional helices not conserved with NS3 (Durr *et al.*, 2005; Liu *et al.*, 2017; *Thoma et al., 2005*). As a result the alignment is restricted to conserved helices. In the case of lobe I and II the RMSD of the fit is 4.9 Å and 6.5 Å respectively (Figure 6 – figure supplement 1A). In the closed state alignment of both domains with the structure of NS3 in the ADP.BeF bound state results in an RMSD of 9.8 Å. Using this alignment the ssDNA bound by NS3 can be docked into the Chd1-Nucleosome structure (Figure 6A). This ssDNA aligns with the top strand of nucleosomal DNA (Figure 6). Conserved motif Ia in ATPase lobe 1 and motifs IV and V from ATPase lobe 2 contact this strand. These residues undergo a ratcheting motion during the course of ATP hydrolysis that drives the ssDNA through NS3 (Gu & Rice, 2010). Similar motion between these residues would be anticipated to drive nucleosomal DNA across the nucleosome dyad in the direction of the longer linker (Figure 6B).

**Figure 6.**
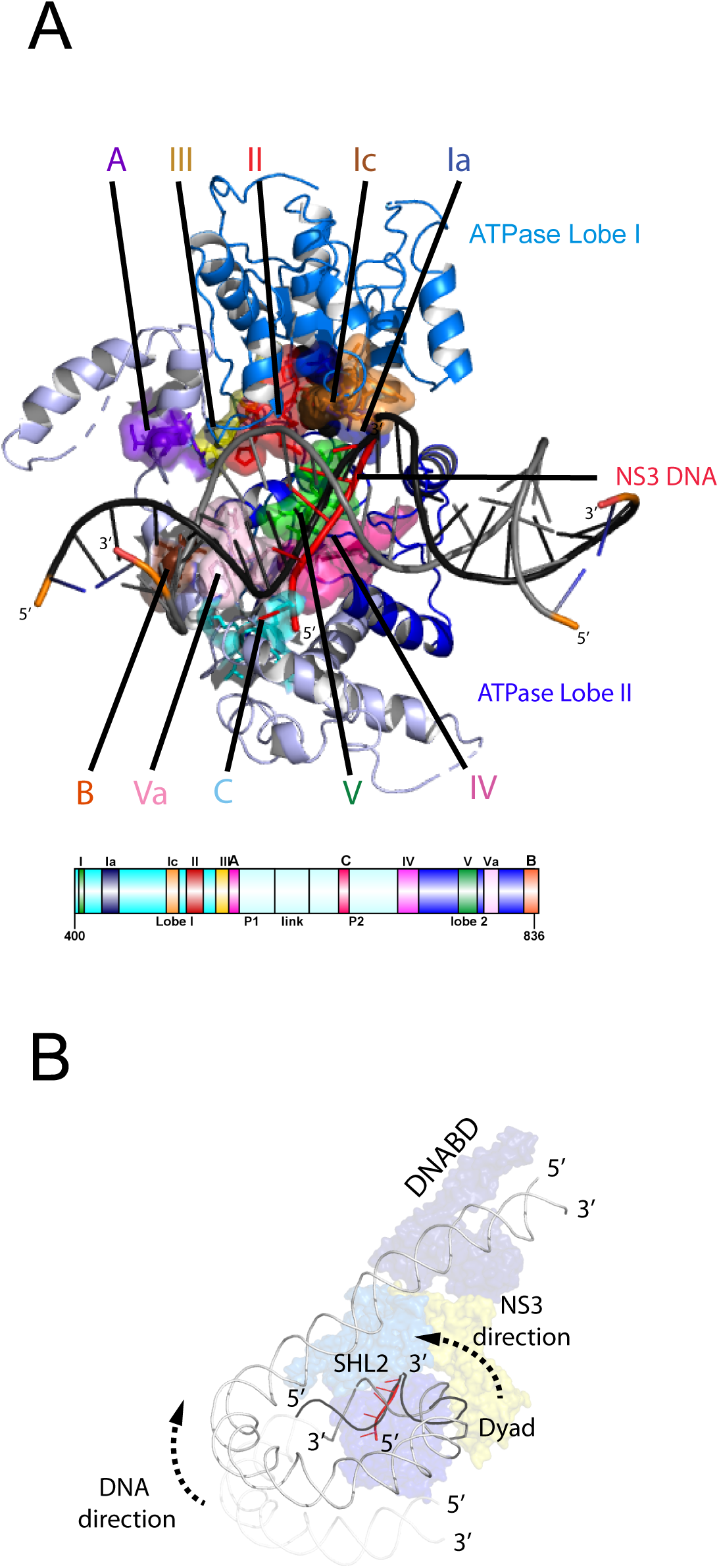
Comparison of NS3 and Chd1 interactions with DNA. **A**) Contacts between motifs conserved with DNA at SHL2. Motifs and colouring are indicated on the structure. The ssDNA from NS3 aligned to Chd1 is shown docked into Chd1 (red). In Chd1 motifs II and III contact the opposite (3’-5’) strand. Contacts made by remodelling enzyme specific extension are labelled A, B and C. The locations of each sequence are indicated in the schematic guide. **B**) Schematic indicating directionality in context of a nucleosome. The directionality of NS3 translocation inferred from docking the ssDNA is 3’-5’ away from the nucleosome dyad. Assuming movement of Chd1 around the nucleosome is constrained (for example via contact with linker DNA, the H4 tail and histone H3) translocation of Chd1 with this directionality is anticipated to drive DNA in the opposite direction towards the long linker as indicated.

It is notable that within Chd1 additional DNA contacts are made that differ from those observed in NS3. Firstly, motifs II and III within lobe 1 contact the opposite DNA strand (Figure 6A). As these motifs are intimately associated with motif Ia they would be anticipated to undergo a similar ratcheting motion with respect to the contacts made by lobe 2. Secondly, the ATPase lobes of Chd1 like those of Snf2 contain additional helices including the protrusions to the helical lobes and the brace helix that are unique to chromatin remodelling enzymes (Figure 6A)(Farnung *et al.*, 2017; Flaus *et al.*, 2006; Liu *et al.*, 2017). These extend the binding cleft between the ATPase lobes and make additional contacts with both DNA strands.

The structure of a fragment of the yeast Snf2 protein bound to a nucleosome revealed contacts between ATPase lobe 1 with DNA at SHL2 and the adjacent DNA gyre at SHL 6 (Liu *et al.*, 2017)(Figure 6 – Figure supplement 2A). The basic surface of lobe 1 responsible for this interaction is not conserved in Chd1, and the acidic residues D464 and E468 make a similar interaction with DNA unlikely. In addition, DNA is not present in this location as it is lifted off the surface of the octamer (Figure 6 – Figure supplement 2B). In the case of the Snf2 protein the interaction with the adjacent DNA gyre is proposed to anchor the translocase preventing it from transiting around the octamer surface (Liu *et al.*, 2017). Chd1 has a relatively small interaction interface with histones, so there is a similar requirement for DNA interactions to constrain motion of the whole protein. In the case of Chd1 this could instead be provided through the interaction of the chromodomains with DNA at SHL1 and through the interaction of the DNA binding domain with linker DNA. Amino acids 476 to 480 of lobe 1 also interact with DNA in the unravelled state (Figure 6 – figure supplement 2B). These residues are not conserved even in Chd1 proteins so the significance of this contact is not clear.

### Two molecules of Chd1 can bind a single nucleosome using the same mode of binding

Chromatin organising motor proteins are capable of catalysing bidirectional nucleosome repositioning that can occur as a result of the binding of two or one enzyme (Blosser *et al.*, 2009; Qiu *et al.*, 2017; Racki *et al.*, 2009; Willhoft *et al.*, 2017). As Chd1 binds to one side of the nucleosome, no steric clashes are anticipated should a second Chd1 bind linker DNA on the opposite side of the nucleosome. To investigate this further, complexes consisting of two Chd1 molecules bound to one nucleosome were prepared using nucleosomal DNA with symmetrical linkers of 14 base pairs and the images processed as indicated (Figure 7 – Figure supplement 1). Most particles were assigned to 2D classes in which two bound Chd1 molecules are discernible, though one is often more dominant likely as a result of the projections of the dominant orientations observed. All 3D classes have two bound Chd1 molecules and the most abundant classes refine to provide an envelope with 15 Å resolution (Figure 7). Two Chd1 molecules bound in the same mode observed in the 1:1 complex can be docked into this volume. There are no direct contacts between the two Chd1 proteins suggesting that the two bound enzymes are likely to function independently. Previously, negative stain EM of two Chd1 molecules bound to a single nucleosome indicated that the DNA binding domain interacted with linker DNA on only one side of the nucleosome (Nodelman *et al.*, 2017). Our envelope shows that both DNA binding domains can bind to linker DNA simultaneously and that the extent of DNA unwrapping is similar on both sides of the nucleosome. This provides further evidence that the differences in DNA binding to the two sides of the 601 nucleosome positioning sequence do not influence the extent of DNA unwrapping. Any differences between the two bound Chd1 molecules must be localised and not detectable at this resolution.

**Figure 7.**
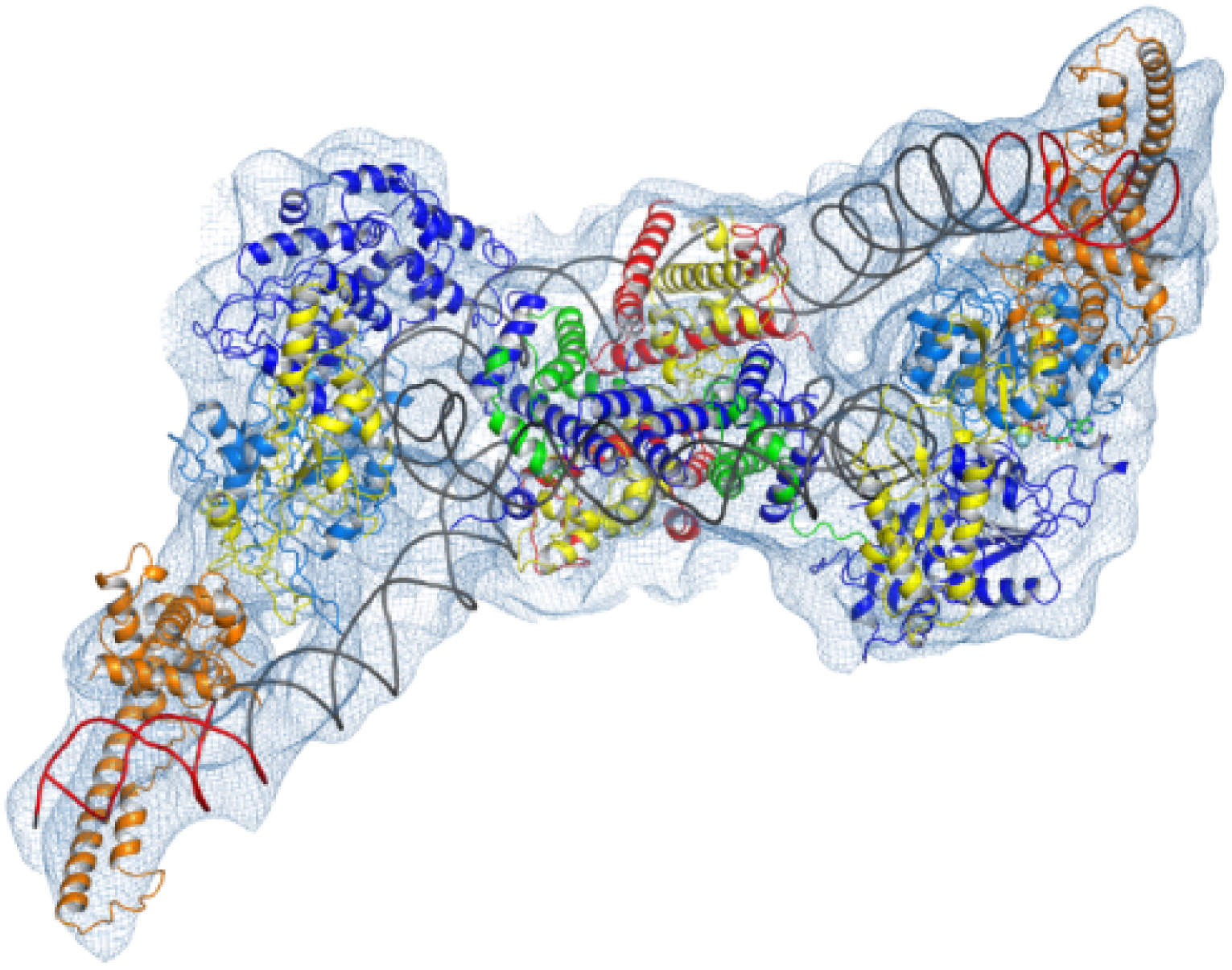
Complex of two Chd1 bound to one nucleosome. Fit of two Chd1 molecules bound to a single nucleosome to density map. The colouring of domains is as for Figure 1.

**Figure 7 – Figure supplement 1.**
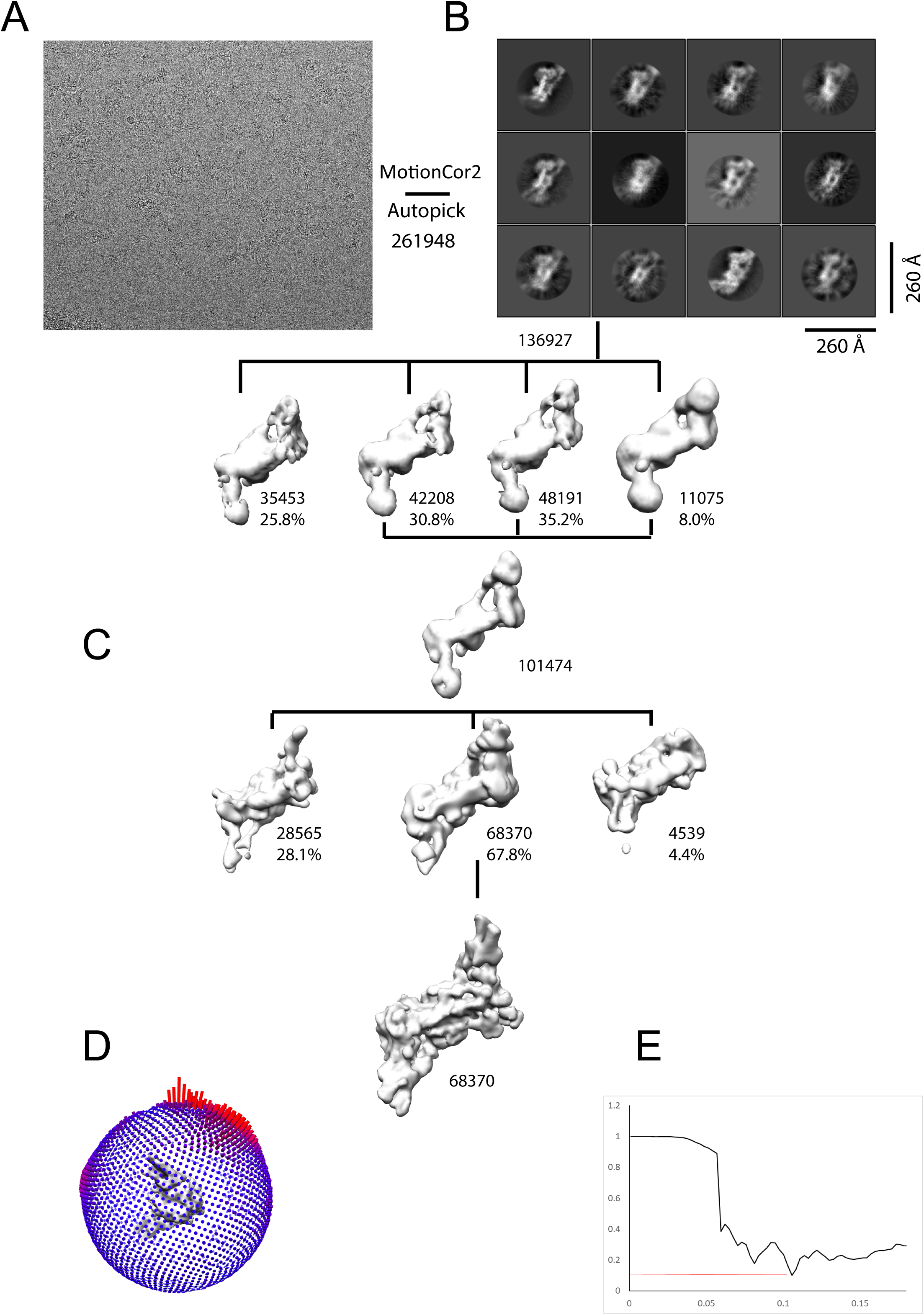
Generation of density map for two Chd1 molecules bound to a single nucleosome. Complexes consisting of two Chd1 molecules bound to a single nucleosome with 14 base pair linkers on each side were prepared on EM grids. **A)** Example micrograph used to identify 261948 particles from which 12 2D classes were identified **B). C)** 3D classes were processed to obtain a final volume based on 68370 particles. Particles display an orientation bias **D),** with a resolution of 16 Å at FSC .143 **E**).

### The trajectory of the histone H3 tail is altered by DNA unwrapping

On the fully wrapped side of the nucleosome the H3 tail can be traced to proline 38, emerging between the DNA gyres at SHL1. In contrast, on the unwrapped side of the nucleosome the H3 tail can be traced to alanine 26 indicating that on this side of the nucleosome the H3 tail is better ordered. In addition, the trajectory of the tail is different to that observed in previous structures (Figure 8A). This altered trajectory was also not observed in a previous structure of a Chd1 bound nucleosome, however, this structure was made in the presence of PAF1 and FACT which may result in some noise in this region that is not apparent in our structure (Farnung *et al.*, 2017). A potential explanation for the defined and altered trajectory of the histone H3 tail on the unwrapped side of the nucleosome is that amino acids within extreme N-terminal region, that are not resolved in our structure, interact with the unravelled DNA. The 25 N-terminal residues include eight lysine and arginine residues that could interact with DNA at different locations along the unravelled linker.

**Figure 8.**
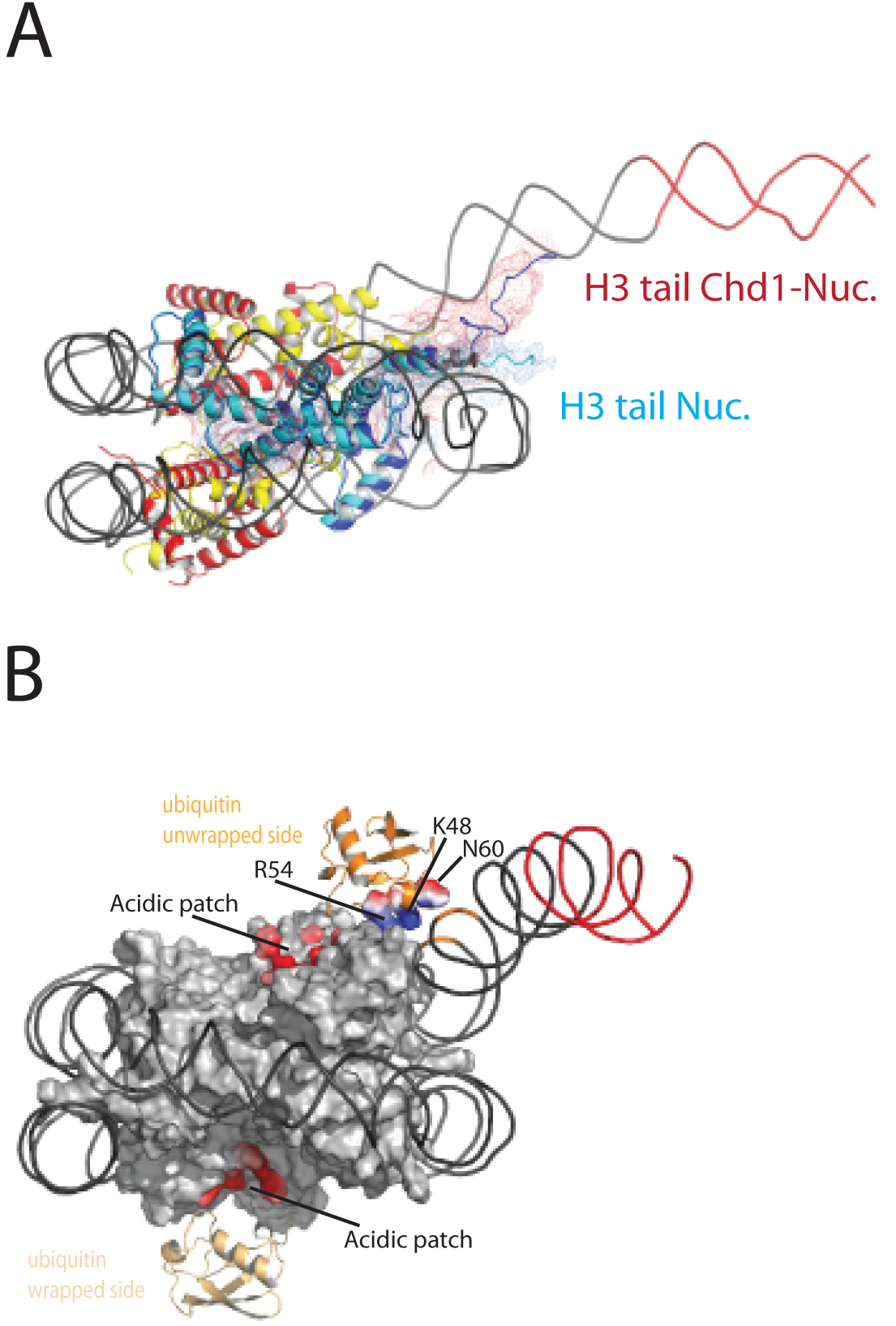
Histone epitopes are repositioned on Chd1 bound nucleosomes. **A)** The histone H3 tail on the unwrapped side of a Chd1 bound nucleosome is shown in dark blue fitted to electron density shown in red. The density extends to residue 26 on this side of the nucleosome and follows a path towards the unwrapped DNA. On the wrapped side of the nucleosome the H3 tail is less well defined and follows a path similar to that observed on nucleosomes bound by 53BP1 (shown in light blue). H3 tail. **B**) Ubiquitin on the wrapped side of the nucleosome is located over the acidic patch. On the unwrapped side of the nucleosome it is repositioned away from the acidic patch and interacts with the unravelled DNA.

### Ubiquitin interacts with unravelled nucleosomal DNA

The electron density for ubiquitin molecules is not as well defined as other components of the complex, and limiting for the overall resolution (Figure 1). This is likely to reflect mobility of the ubiquitin peptides. Consistent with this the electron density determined from X-ray diffraction on crystallised nucleosomes with ubiquitin conjugated at this location resulted in no density attributable to ubiquitin (Machida *et al.*, 2016). On the wrapped side of the nucleosome, ubiquitin is located adjacent to the acidic patch that is widely used as an interface for nucleosome binding proteins (McGinty & Tan, 2016) (Figure 8B). This is also the location that ubiquitin conjugated to H2A K15 has been observed to occupy on unbound nucleosomes (Wilson *et al.*, 2016) and likely represents a favourable conformation for ubiquitin when coupled at different sites within this locality (Vlaming *et al.*, 2014).

On the unwrapped side of the nucleosome, the ubiquitin peptide is displaced across the lateral surface towards the DNA. The unwrapped DNA is oriented away from the lateral surface and this positions the DNA backbone at SHL6 in contact with the repositioned ubiquitin (Figure 8B). In 2D classes, ubiquitin is more prominent on the unwrapped side suggesting it is more tightly constrained. The residues interacting with DNA include Lys 48 Arg 54 and Asp 60. It is likely this interaction stabilises DNA in the unwrapped state, perhaps explaining why H2B K123 ubiquitinylation directly stimulates remodelling by Chd1(Levendosky *et al.*, 2016).

## Discussion

A striking feature of the Chd1 nucleosome complex is the limited number of contacts with histones. The two direct contacts with histones are with the H3 alpha one helix and with the histone H4 tail (Figure 2 – Figure supplement 4). The contact is with the H4 tail is required for efficient remodelling by Chd1 (Ferreira *et al.*, 2007) as it is for ISWI subfamily enzymes (Clapier *et al.*, 2001; Hamiche *et al.*, 2001). Aside from these two contacts, the interaction of Chd1 with nucleosomes is dominated by interactions with DNA. This leaves the majority of the nucleosome accessible for binding by additional factors. We show that a second Chd1 molecule can bind the opposite side of the nucleosome using a similar mode of interaction (Figure 7). Even in the case of a nucleosome bound by two Chd1 molecules the lateral surfaces of the nucleosome are accessible for binding by other factors.

Despite a lack of direct contacts with histones H2A and H2B, Chd1 activity is dependent on histone dimers (Levendosky *et al.*, 2016). The requirement for histone dimers may arise as a result of dimer loss affecting DNA wrapping. Loss of a histone dimer will result in a loss of histone DNA contacts at SHL3.5, 4.5 and 5.5. More extensive unwrapping of DNA to SHL3.5 would require major repositioning of DNABD in order to retain the interaction with linker DNA while the chromoATPase is engaged at SHL-2. This provides a potential explanation for the dependency of Chd1activity on histone dimers. Conversely, association of a histone dimer with a histone tetramer or hexamer around which DNA is initially significantly unwrapped, could be stabilised by the binding of Chd1 rewrapping DNA to SHL5. The stabilisation of the SHL5 wrapped state may facilitated correct docking of histone dimers into chromatin. This provides a mechanistic basis for the observed activities of Chd1 in H2A/H2B transfer (Lusser *et al.*, 2005) and chromatin assembly (Fei *et al.*, 2015; Lee *et al.*, 2012; Torigoe *et al.*, 2013). Repositioning of the Chd1 DNA binding domain towards the major orientation observed in free solution, the Apo state reported by (Sundaramoorthy *et al.*, 2017), would guide linker DNA towards the fully wrapped state. As a result Chd1 has the potential to function in multiple stages of chromatin assembly and the generation of organised chromatin (Lusser *et al.*, 2005; Robinson & Schultz, 2003; Torigoe *et al.*, 2013).

The Chd1 enzyme has the ability to organise spaced arrays of nucleosomes both in vitro and in vivo (Gkikopoulos *et al.*, 2011; Lusser *et al.*, 2005; Robinson & Schultz, 2003). Enzymes that exhibit this organising activity typically reposition nucleosomes away from the ends of short DNA fragments. This is also true for Chd1 (McKnight *et al.*, 2011; Stockdale *et al.*, 2006). As a result we would anticipate that Chd1 would be most likely to reposition nucleosomes away from the short (exit) linker, encroaching into the long (entry) linker. Repositioning of nucleosomes with this directionality conflicts with the directionality of translocation inferred from docking the tracking strand of NS3 into Chd1(Farnung *et al.*, 2017)(Figure 6). Tracking along this strand with 3’-5’ directionality would instead be anticipated to draw DNA into the nucleosome from the exit side.

Inferring the mechanism of Chd1 from NS3 is complicated by the fact these enzymes are not so closely related. Conserved motifs are difficult to align based on sequence alone. In addition some aspects of nucleic acid binding by both Snf2 and Chd1 profoundly differ from NS3. Notably, motifs II and III within lobe 1 contact the opposite, 3’-5’ stand, which is not present in NS3. In addition, Snf2 related chromatin remodelling enzymes contain features that extend the nucleic acid binding cleft between the two ATPase lobes and make contacts with both strands (Figure 6). As a result of the extensive contacts with both strands, it is possible that the assignment of guide and tracking strands within remodelling ATPases is not absolute as tracking may be coupled to both strands. Consistent with this experiments that have probed the action of remodelling enzymes using short gaps in either strand of nucleosomal DNA have found them to be sensitive to lesions in either strand (Saha *et al.*, 2005; Zofall *et al.*, 2006).

The introduction of gaps in nucleosomal DNA has also been used to infer the directionality with which ATPases’s move along DNA. Introduction of gaps distal to the SHL 2 location closest to the entry linker DNA has been observed to impede the action of Snf2, Iswi and Chd1 enzymes (McKnight *et al.*, 2011; Saha *et al.*, 2005; Zofall *et al.*, 2006). This has been used as evidence that the enzyme translocation that drives repositioning initiates from the SHL 2 located distal to the entry DNA. As a consequence it has been proposed that Chd1 bound in the cross gyres conformation, that we and others observe, represents an inactive state in which the interaction of the DNABD with linker DNA is inhibitory (Farnung *et al.*, 2017; Nodelman *et al.*, 2017). If this were the case, then Chd1 directed repositioning might be expected to be limited once a nucleosome has moved far enough from a DNA end to enable the DNABD to contact linker DNA. On Chd1 bound nucleosomes the DNABD contacts extend to 7bp from the nucleosomes edge. A stall to repositioning after 7bp is not observed, instead Chd1 repositions nucleosomes 23-39 base pairs into a 54 base pair linker in a very similar way to Isw1a and Isw2 (Stockdale *et al.*, 2006).

There is further evidence to support the conformation of Chd1 with the DNABD bound to linker DNA as an active state. Firstly, in the ADP.BeF bound state, the ATPase domains are repositioned to a closed conformation and conserved residues are positioned for catalysis (Figure 2 – figure supplement 3). The closure of the ATPase domains in the Chd1-nucleosome complex is connected to the nucleotide-dependent unwrapping of DNA via repositioning of the chromodomains, which in turn levers the DNABD position. Secondly, efficient nucleosome repositioning by Chd1 is dependent on the DNABD (Ryan *et al.*, 2011b). This is anticipated if the DNA binding domain acts to generate an active conformation, but not if sensing of exit linker DNA is repressive. Thirdly, the activity of chimeric Chd1 proteins in which DNA binding is provided via a heterologous domain is greatest when the cognate binding site is placed in the entry linker mimicking the arrangement observed in the Chd1-nuceosome structure (McKnight *et al.*, 2011; Patel *et al.*, 2012). An additional confounding factor in assigning the directionality with which Chd1 translocates is the recent observation that a single Chd1 molecule can direct bidirectional nucleosome movement (Qiu *et al.*). As a result, further studies are required to resolve how Chd1 acts to drive DNA across the octamer surface.

Both budding yeast Chd1 and human Chd2 are found to be enriched within coding regions (de Dieuleveult *et al.*, 2016; Gkikopoulos *et al.*, 2011; *Lee et al., 2017*). Histone H3 K36me3 is a hallmark of coding region nucleosomes, so we prepared nucleosomes alkylated to mimic trimethylation at this position. Alkylation modestly stimulates Chd1 activity (Figure 1 – Figure supplement 1), raising the possibility that this modification is recognised by the enzyme, possibly via the chromodomains. However, we observe electron density for the histone H3 tail to residue 26, indicating that H3K36 does not stably interact with the chromodomains or any other component of Chd1 in the structure reported here. Furthermore, for this interaction to occur, either the chromodomains would need to be repositioned, or the structure of the N-terminus of H3 reconfigured for example by unfolding of the alpha–N helix (Elsasser *et al.*, 2012; Liu *et al.*, 2012).

The improved density for the H3 tail on the unwrapped side of the nucleosome is most likely to result from the interaction of the basic N-terminal region of the H3 tail, which is not resolved, with DNA. It is notable that in the fully wrapped state the H3 tail would need to follow a very different path in order to interact with DNA. Consistent with this the trajectory of the H3 tail on the unwrapped side of the nucleosome is different to that observed in structures of intact nucleosomes (Wilson *et al.*, 2016). This raises the possibility that changes to DNA wrapping could affect the way in which histone tail epitopes are displayed. In principle, such effects could be positive or negative. For example the tudor domain of PHF1 preferentially interacts with trimethylated H3K36 on partially unwrapped nucleosomes (Gibson *et al.*, 2017). The interaction of the PHD domains of Chd4 with DNA is also inhibited by nucleosomal DNA (Gatchalian *et al.*, 2017). As a result if Chd4 generates unwrapped structures similar to those observed with Chd1 the interaction of these domains would be enhanced. The reconfiguration of the H3 tail by Chd1 has the potential to affect the interaction of histone reader, writer and eraser enzymes with the tail and as a result the distribution of these modifications in chromatin. Such effects have been observed, as Chd1 contributes to the establishment of boundaries between H3K4me3 and H3K36me3 at most transcribed genes (Lee *et al.*, 2017).

H2BK120 ubiquitination is also enriched in coding region chromatin, and has previously been observed to stimulate Chd1 activity (Levendosky *et al.*, 2016). Although the level of Chd1 stimulation is only 2-fold, we have previously observed that mutations exerting a 2-fold effect on Chd1 activity *in vitro* affect nucleosome organisation *in vivo* (Sundaramoorthy *et al.*, 2017). It has also been observed that H2BK120Ub negatively affects the activity of some ISWI containing enzymes (Fierz *et al.*, 2011). As organisation of coding region nucleosomes involves these and other enzymes (Krietenstein *et al.*, 2016; Ocampo *et al.*, 2016; Parnell *et al.*, 2015), H2BK120Ub has the potential to regulate interplay between different enzymes.

Ubiquitin on the unwrapped side of the nucleosome is repositioned such that it interacts directly with DNA. As in the case of the H3 tail, the repositioning of the ubiquitin resulting from Chd1-directed DNA unwrapping could potentially affect interactions with the factors involved in the placing, removal or recognition of H2BK120ub. The most striking functional evidence for this interplay is that H2BK120ub is greatly reduced in Chd1 mutants (Lee *et al.*, 2012). One possible explanation for this effect is that Chd1 sequesters ubiquitin in a conformation less accessible for removal. Consistent with this the position of ubiquitin on the unwrapped side of Chd1 bound nucleosomes is incompatible with interaction with the SAGA DUB module (Morgan *et al.*, 2016). Interestingly, the paradigm for trans regulation between histone modifications stems from the interplay between H2BK120ub and H3 K4 methylation (Sun & Allis, 2002), both of which are influenced by Chd1 binding. While Chd1 is not required for H3K4me3 (Lee *et al.*, 2012) it does influence the distribution of this histone modification (Lee *et al.*, 2017).

H2BK120Ub has previously been observed to directly affect chromosome structure at the level of chromatin fibre formation (Debelouchina *et al.*, 2017; Fierz *et al.*, 2011). Our observations show a new role for H2BK120Ub at the level of nucleosomal DNA wrapping. The specific relocation of ubiquitin on the unravelled side of the nucleosome, the local distortion of H2B at the site of attachment and the presence of lysine and arginine residues at the site of interaction with DNA all indicate this is a favourable interaction that stabilises DNA in the unwrapped state. The outer turns of nucleosomal DNA rapidly associate and dissociate on millisecond time scales, with occupancy of the unwrapped state estimated at 10% (Li *et al.*, 2005). The ubiquitin interaction we have observed would be anticipated to stabilise the transiently unwrapped state increasing its abundance. It is however, unlikely that the unwrapped state predominates in the absence of Chd1 or other factors that promote unwrapping as the structure of isolated ubiquitinylated nucleosomes is unchanged (Machida *et al.*, 2016). Nonetheless, increased occupancy of the transiently unwrapped state would be anticipated to facilitate access to nucleosomal DNA. Chromatin folding to form higher order structures is likely to be favoured by fully wrapped, nucleosomes and so an increase in the proportion of unwrapped nucleosomes could potentially contribute to the effects of H2BK120Ub on chromatin fibre formation (Fierz *et al.*, 2011). Many other processes involving chromatin dynamics are linked to H2BK120Ub including transcription (Bonnet *et al.*, 2014), DNA repair (Moyal *et al.*, 2011; Nakamura *et al.*, 2011) and DNA replication (Lin *et al.*, 2014). A more stable unwrapped state could also provide an explanation for the association of factors that lack recognised ubiquitin interaction domains, with ubiquitinylated chromatin (Shema-Yaacoby *et al.*, 2013). Interestingly, H2BK120Ub associating proteins include human Chd1, SWI/SNF complex, pol II and the elongation factors NELF and DISF (Shema-Yaacoby *et al.*, 2013).

The change in the position of ubiquitin also has the potential to indirectly affect the way in which other factors interact with ubiquitinylated nucleosomes. On the wrapped side of the nucleosome ubiquitin is positioned such that it occludes access to the acidic patch formed by the cleft between histones H2A and H2B. This provides surface via which many proteins including LANA peptides (Barbera *et al.*, 2006), RCC1 (Makde *et al.*, 2010), Sir3 (Armache *et al.*, 2011), PRC1 (McGinty *et al.*, 2014) and the SAGA DUB module (Morgan *et al.*, 2016) interact with nucleosomes. The repositioning of ubiquitin away from the acidic patch on the unwrapped side of the nucleosome improves access to the acidic patch. In this way H2BK120ub may provide a means of regulating access to the acidic patch that is sensitive to changes in nucleosome structure.

Although the repositioning of the H3 tail and ubiquitin were observed on Chd1 bound nucleosomes, the potential for reconfiguration of histone epitopes may be more general. All processes that generate local DNA unwrapping would be anticipated to result in similar repositioning of histone tail epitopes. In particular, where combinations of modifications are recognised bivalently, the spatial alignment of epitopes will be important for recognition by coupled reader domains. This potentially provides a means of tuning signalling via histone modifications to regions of transient histone dynamics.

## Acknowledgements

We thank Daniel Claire electron Bio-Imaging Centre (eBIC), Diamond light source Ltd, UK for data collection with respect to the 4.5 Å structure. We thank Rebecca Thompson and Neil Ranson for assistance with collection of 2Chd1 bound to one nucleosome structure at the Astbury Biostructre Laboratory, University of Leeds. The ubiquitin expression plasmid was kindly provided by Ron Hay, University of Dundee. This work was funded by Wellcome Senior Fellowship 095062, Wellcome Trust grants 094090, 099149 and 097945. ALH was funded by and EMBO long term fellowship ALTF 380-2015 co-funded by the European Commission (LTFCOFUND2013, GA-2013-609409). We Thank Dale Wigley and Chris Aylett for useful discussions.

## Methods

### Cloning, protein expression and purification

ScChd1 C-terminal and N-terminal truncations were made from the full length clone described in Ryan et al, using an inverse PCR strategy (Ryan *et al.*, 2011a). Site directed mutagenesis was used to introduce cysteine residues at strategic locations on ScChd1 1-1305ΔC using standard cloning procedure. All proteins were expressed in Rosetta2 (DE3) pLysS Escherichia Coli cells at 20° C in Auto-induction media and the purification of the protein was carried out typically as described in Ryan et al. After the purification of the protein the GST tag was cleaved with precision protease and the tag cleaved proteins were subjected to size exclusion chromatography using Superdex S200 10/300 GL columns (GE Healthcare). Expression and purification of Xenopus laevis histones were carried out as described previously (Luger *et al.*, 1999).

### Installation of Methyl-lysine analogues in H3 K36

Alkylation of cysteine-mutant histones to generate histones modified with methyl-lysine analogues was performed as in (Simon M.D. et al, Cell 2007). Approximately 10mg of lyophilised cysteine mutant histone was resuspended in 800uL (me3) or 900uL (me0) degassed alkylation buffer (1M HEPES, 10mM D,L-methionine, 4M Guanidine HCl, pH7.8). Histones were reduced with fresh 30mM DTT for 30 minutes at room temperature.

For trimethyl-lysine analogues, the reduced histone was added to approximately 125mg of (2-Bromoethyl) trimethylammonium bromide (Sigma 117196-25G) in 200uL of DMF and incubated in the dark at 50⁰C for 3 hours. An additional 10uL of DTT was added, and the reaction was allowed to proceed overnight at room temperature.

For generation of the unmethylated lysine analogue, 75uL of 1M 2-Bromoethylamine hydrobromide (Fluka 06670-100G) was added to the reduced histone and was incubated at room temperature in the dark for 3 hours. An additional 10uL of DTT was added for 30 minutes prior to the addition of an extra 75uL of alkylating agent, and the reaction was allowed to proceed overnight at room temperature in the dark.

The reaction was terminated with the addition of 50uL 2-mercaptoethanol for 30 minutes and the alkylated histone was desalted either by dialysis into water with 2mM 2-mercaptoethanol or on a PD-10 desalting column (GE 52130800). The shift in molecular weight associated was confirmed via MALDI-TOF mass spectrometry.

### In vitro ubiquitination

Recombinant expression of xH2B K120C and His-TEV-Ubiquitin G76C mutant proteins was induced with IPTG for 4 hours in Rosetta 2 DE3 pLysS cells grown at 37⁰C. Inclusion body purification followed by cation exchange chromatography was performed to isolate the histone protein. Ubiquitin was purified using HisPur cobalt resin with 150mM sodium chloride/20mM Tris pH8 buffer and eluted with 350mM imidazole. Histones and ubiquitin were desalted by dialysis into water with 2mM 2-mercaptoethanol and lyophilised for storage. Proteins were re-suspended in 50mM ammonium bicarbonate pH 8 and treated with 2mM TCEP for 1 hour. Ellman’s reagent was used to ascertain the concentration of free sulfhydryls, and xH2b and ubiquitin were combined at equimolar ratios, as defined by the Ellman’s assay, and diluted with 50mM ammonium bicarbonate to 200-250uM each protein. The proteins were crosslinked at room temperature with four hourly additions of ¾ molar ratio of 1,3 dichloroacetone (freshly prepared in DMF). An equal volume of denaturing buffer (7M Guanidine HCl, 350mM sodium chloride, 25mM Tris pH7.5) was added to the reactions, which were purified using HisPur cobalt resin, pre-equilibrated in denaturing buffer. The His-TEV-Ub-xH2B crosslinked product was eluted with 350mM imidazole and dialysed into SAUDE200 buffer (7M Urea, 20mM sodium acetate, 200mM sodium chloride, 1mM EDTA, 5mM 2-mercaptoethanol) overnight. The ubiquitinated histone was further purified over a cation exchange column, as before, and fractions were dialysed into water with 2mM 2-mercaptoethanol and lyophilised for storage.

### Preparation of recombinant nucleosomes

Xenopus H2B-K120 ubiquitinylated histones were refolded in equimolar ratios with H2A and similarly H3 K36 methyl analogue histones were refolded in equimolar ratios with histone H4 to obtain dimers and tetramers as described previously for wild type histones Dyer et al., and purified on a size exclusion chromatography using S200 gel filtation column. The peak fractions were analysed with SDS-PAGE gel and pooled. 601 DNA fragments of respective lengths for recombinant nucleosome assembly were generated by PCR method as described previously (Sundaramoorthy *et al.*, 2017). Nucleosomes were generated by salt dialysis as described previously by combining H2A/H2B-K120 ubiquitin dimer, H3K36 methyl lysine analogue tetramer (2:1 ratio) with DNA containing PCR amplified Widom 601 DNA sequence.

### Nucleosome Repositioning Assay

Nucleosomes were reconstituted on Cy3 (me0) and Cy5 (me3) labelled DNA, based on the 601 sequence, with a 47bp extension. Repositioning by Chd1 was performed in 40mM Tris pH7.4, 50mM KCl, 3mM MgCl2, 1mM ATP, 100nM each nucleosome, and 10nM Chd1; 10uL was removed at each time point (T=0, 4, 8, 16, 32, and 64 minutes), placed on ice, and stopped with the addition of 100ng/uL competitor DNA, 200mM NaCl, and 1.6% sucrose. Repositioned nucleosomes were run on 6% PAGE/0.2X TBE gels in recirculating 0.2X TBE buffer for 3-4 hours at 300V. The percent of repositioned nucleosomes was analysed using Aida image analysis software. Data were fit to a hyperbola in Sigma Plot, to determine the initial rate of repositioning.

### Nucleosome binding

*Xenopus laevis* nucleosomes (20nM), reconstituted on Cy3 labelled 0W11 DNA, were bound to titrations of Chd1 enzymes (concentration specified in figure legend) in 50mM Tris pH 7.5, 50mM sodium chloride, and 3mM magnesium chloride supplemented with 100ug/mL BSA. Unbound and bound nucleosomes were separated on a pre-run 6% polyacrylamide gel (49:1 acrylamide: bis-acrylamide) in 0.5X TBE buffer for 1 hour at 150V. The gel shift was scanned on Fujifilm FLA-5100 imaging system at 532nm.

### Spin labelling of ScChd1, PELDOR measurements and modelling

MTSL was conjugated to introduce cysteines immediately following size exclusion purification as described in Hammond et al (Hammond et al., 2016). Excess unreacted labels were removed from the sample by dialysis. PELDOR experiments were conducted at Q-band (34 GHz) operating on a Bruker ELEXSYS E580 spectrometer with a probe head supporting a cylindrical resonator ER 5106QT-2w and a Bruker 400 U second microwave source unit as described previously (Hammond et al., 2016). All measurements reported here were made at 50K. Data analysis was carried out using the DeerAnalysis 2013 package (Jeschke and Polyhach, 2007). The dipolar coupling evolution data were first corrected to remove background decay. Tikhonov regularisation was then used to determine distance distributions from each dataset.

To model the distance distribution for the open conformation of Chd1 helicase lobes crystal structure of chromo helicase (PDB Code: 3MWY) (Hauk et al., 2010) was used. For the closed conformation refined cryoEM structure of Chd1 bound to nucleosome in the presence of ADP.BeF_x_ described in this study was used as a model. For each structure, R1 spin labels were added and the distribution simulated for each position using MTSSL wizard in Pymol. Also the average distance from the distribution from a pair of spin labels were calculated using MTSSL wizard in Pymol.

### Sample preparation, Cryo Electron Microscopy data collection and analysis

The appropriate ratio of ScChd1(1-1305Δ57-88) to nucleosome for 1:1 and the 2:1 complex formation in the presence of 5-fold molar excess of ADP-BeF_x_ was determined by titration and native PAGE analysis. The formed complex was then purified by size exclusion gel filtration using a PC 3.2/30 superdex 200 analytical column in 20mM Tris, 50mM NaCl and 250 µM ADP.BeF_x_. In a typical run 50uLs of 20 µM of sample was injected using Dionex autoloader. 50uLs fractions were collected and analysed in 6% Native PAGE gel and appropriate fractions containing ScChd1-nucleosome complexes were pooled together. A 4 µl drop of sample was then applied to C-flat Holey carbon foil (400 mesh R1.2/1.3 uM) pre-cleaned with glow discharge (Quorum technologies). After 15 second incubation, grids were double side blotted for 4 s in a FEI cryo-plunger (FEI Mark III) at 90% humidity and plunge frozen into −172 °C liquefied ethane. Standard vitrobot filter paper Ø 55/20 mm, Grade 595 was used for blotting.

The prepared grids are initially checked for its ice quality and the particle distribution using a JEOL 2010 microscope with side-entry cryo-holder operated at 200 keV and equipped with a gatan 4k × 4k CCD camera. Cryo-grids were then stored in liquid nitrogen and dry-shipped to respective centre for data collection. For the 1:1 complex the data was acquired on a FEI Titan Krios transmission electron microscope (TEM) operated at 300 keV, equipped with a K2 summit direct detector (Gatan). Automated data acquisition was carried out using FEI EPU software at a nominal magnification of 105,000×.

For the 2:1 complex the data was collected at Astbury centre for cryo electron microscopy, Leeds, on a FEI Titan Krios transmission electron microscope (TEM) operated at 300 keV, equipped with Falcon 3 detector (FEI).

The movie frames were subjected to frame wise motion correction using MotionCor2. CTF correction was then performed on the motion corrected summed image using Gctf. Subsequent image processing was performed with RELION 2.0.4. About 5000 particles from 50 micrographs were first handpicked in RELION, extracted and 2D classes were generated. These 2D classes were then used as a reference in RELION autopick routine and particles were picked from respective number of micrographs from each dataset. The autopicked particles were subsequently extracted and sorted. An iterative round of two-dimensional classification was performed to discard poorly averaging particles, contamination and exploded particles. On the resultant cleaned up particle stack a hierarchical three-dimensional classification and refinement was performed as described in the results. A low pass filtered low resolution chd1 engaged nucleosome structure was used as an initial model in the 3D classification. At the three-dimensional refinement stages a mask that encompasses the entire Chd1-nucleosome complex was applied. Post processing of refined models was performed with automatic B factor determination in RELION. Local resolution estimates were determined using Resmap-1.4.

## Model Building

For model building X.laevis nucleosome with Widom 601 sequence (PDB 3LZ0), the S.cerevisiae Chd1 DNA-binding domain (PDB 3TED) and the ATPase core with tandem chromo domain (3MWY) were used. The domains were individually placed into the electron density using UCSF chimera and fitted as a rigid body. The path of the unwrapped DNA, H4 tail region and H3 tail region were manually built in Coot. Protein back bone restraints and DNA base pair, and parallel pair restraints were generated using ProSMART and LibG modules. The generated restraints were then used as constraint in jelly body refinement with CCPEM REFMAC program. ADP⋅BeF_3_ was built by superpositioning ATP-gamma-S from the inactive Chd1 structure (PDB code 3MWY) onto our model, inspected in COOT and replacing the ATP analogue with ADP⋅BeF_x_. Sequence alignments were generated using JALVIEW (Waterhouse *et al.*, 2009). The EM volumes and the fitted model can be accessed from the EMDB database with EMDB accession number EMD 4336; PDB 6G0L EMD-4318 and PDB code 6FTX.

---

**Figure 6 – Figure supplement 1. Comparison of NS3 and Chd1 ATPase domains.** ATPase domains of Chd1 and NS3 docked individually and together. ATPase lobes 1 and 2 are shown in light and dark blue. Regions specific for Snf2 proteins are shown in white. NS3 ATPase domains in raspberry.

**Figure 6 – Figure supplement 2. Snf2 and Chd1 interactions with the adjacent DNA gyre. A**) Contacts between Snf2 and DNA adjacent to SHL2 at SHL6 K855, R880 and K885 (PDB 5X0Y). **B)** These contacts are not conserved in Chd1 and instead an acidic surface is presented by D464 and E468. The region 476 to 480 contacts the unravelled DNA. In both panels normally wrapped nucleosomal DNA is indicated in orange, and the gyre unwrapped by Chd1 in black.

